# Developing epidemiological preparedness for a plant disease invasion: modelling citrus huánglóngbìng in the European Union

**DOI:** 10.1101/2024.06.04.597414

**Authors:** John Ellis, Elena Lázaro, Beatriz Duarte, Tomás Magalhães, Amílcar Duarte, Jacinto Benhadi-Marín, José Alberto Pereira, Antonio Vicent, Stephen Parnell, Nik J. Cunniffe

## Abstract

Huánglóngbíng (HLB; citrus greening) is the most damaging disease of citrus worldwide. While citrus production in the USA and Brazil have been affected for decades, HLB has not been detected in the European Union (EU). However, psyllid vectors have already invaded and spread in Portugal and Spain, and in 2023 the psyllid species known to vector HLB in the Americas was first reported within the EU. We develop a landscape-scale, epidemiological model, accounting for heterogeneous citrus cultivation and vector dispersal, as well as climate and disease management. We use our model to predict HLB dynamics following introduction into high-density citrus areas in Spain, assessing detection and control strategies. Even with significant visual surveillance, we predict any epidemic will be widespread on first detection, with eradication unlikely. Introducing increased inspection and roguing following first detection, particularly if coupled with intensive insecticide use, could potentially sustain citrus production for some time. However, this may require chemical application rates that are not permissible in the EU. Disease management strategies targeting asymptomatic infection will likely lead to more successful outcomes. Our work highlights modelling as a key component of developing epidemiological preparedness for a pathogen invasion that is, at least somewhat, predictable in advance.

## Introduction

Consequences of plant disease epidemics threaten ecosystem services (Boyd et al., 2013) and food security (Strange and Scott, 2005). Emerging pathogens, which cause disease in new locations or on new plant host species, can be particularly damaging (Ristaino et al., 2021). However, emerging epidemics are well documented (Rosace et al., 2023; Jeger et al., 2023; Fielder et al., 2024), and invasion rates are increasing (Bebber et al., 2014). Drivers include changes to farming practices and land use (Anderson et al., 2004), climate change (Singh et al., 2023), and increased travel and trade (Brasier, 2008).

Rising invasion rates have focused attention on how emerging epidemics can be detected and controlled (Cunniffe and Gilligan, 2020). It is particularly important to anticipate and be able to react quickly to invasions, since this gives control the best chance of success (Fraser et al., 2004). But effective detection and control strategies can be hard to devise for invading pathogens because epidemiology in new locations is inadequately characterised (Thompson et al., 2018). Mathematical modelling can play a key role. Models offer a rational basis to integrate what is known with what is not known to design surveillance (Parnell et al., 2017) and to determine when, where and how to control disease (Cunniffe et al., 2015a). However, modelling of emerging plant pathogens has very often been done retrospectively (e.g., Cunniffe et al. (2016), Radici et al. (2024)).

Here we focus on modelling in advance of an invasion that is, at least somewhat, predictable. We use citrus greening (aka huánglóng-bíng, HLB) in the European Union (EU) as a timely and socioeconomically important case study. Worldwide, citrus is an important crop, and HLB its most devastating disease (Gottwald, 2010). HLB has been reported in over 60 countries (Zhang et al., 2023), and impacts on the citrus industries of Brazil and the USA are significant. For example, since 2005 citrus production in Florida has decreased by 80%, whereas in Brazil over 64 million trees have been removed in São Paulo state (Graham et al., 2024). However, HLB has not been reported in the EU, and citrus production in the Mediterranean Basin remains unaffected (Wang, 2020), although recent discoveries of a high-profile vector species in Israel (EPPO, 2022) then in the EU itself in Cyprus (EPPO, 2023) are concerning.

HLB is associated with three non-cultivable phloem-restricted bacteria: *Candidatus* Liberibacter asiaticus (*C*Las), *Ca* L. africanus (*C*Laf) and *Ca* L. americanus (*C*Lam) (Bové, 2006). HLB is primarily transmitted by two insect vectors, the Asian citrus psyllid (ACP), *Diaphorina citri* Kuwayama (Hemiptera: Psyllidae), and the African citrus psyllid (AfCP), *Trioza erytreae* Del Guercio (1918) (Hemiptera: Triozidae). ACP has been found in Asia, North America, South America, and a few locations in Africa. There have also been recent detections of ACP in Israel (EPPO, 2022) and Cyprus (EPPO, 2023). AfCP has been found in many countries in Africa and the Middle East, and was detected in Europe in 2014 (Perez-Otero et al., 2015), with subsequent spread over large areas of northwestern Spain and western Portugal (EFSA et al., 2019b). It has been shown experimentally that both ACP and AfCP vector *C*Las (Reynaud et al., 2022), the most aggressive of the three bacteria causing HLB (Bové, 2006).

Both the recent detections of ACP in Israel and Cyprus and the established presence of AfCP in Spain and Portugal are concerning, since either vector could facilitate HLB transmission were a pathogen to be introduced (Cocuzza et al., 2017). Although contingency plans for the arrival of HLB in Spain and Portugal exist (DGAV, 2021; BOE, 2023), and have been tested in formal simulation exercises (Aragón et al., 2022), designing an effective response is challenging. Management interventions that have been used somewhat successfully in other areas might not translate to the EU, since important epidemiological aspects are different. For example, there are differences in how commercial and non-commercial citrus are distributed (Moreira et al., 2019), in climatic drivers of vector population dynamics (Cocuzza et al., 2017), and in regulations dictating which pesticides can be applied and how often (Urbaneja et al., 2020). Each of these factors has knock-on effects upon outbreak management. This is precisely when modelling is most useful.

Here we show how modelling can contribute to developing epidemiological preparedness for a plant pathogen. We have developed a flexible and transferable stochastic landscape-scale model, which accounts for heterogeneity in the citrus host landscape, spatial spread of a vector (including effects of climate and disease management on its population dynamics), and the concomitant spread of HLB. We focus here on the Iberian Peninsula – Spain and Portugal – driven by the availability of citrus density data, and the status of Spain as the largest citrus producer in the EU (Schimmenti et al., 2013). Data from the spread of AfCP in Spain and Portugal to date is used to parameterise psyllid dispersal in our model. However, the fitted model can be applied to any EU region, assuming climatic and citrus host data were available.

While our model must track spread across large areas of Spain and Portugal when fitting psyllid dispersal parameters, our major focus is to assess and compare surveillance and control strategies before and during the early stage of any epidemic. We do not aim to predict precisely where in the EU the pathogen will enter, and so do not attempt to model relative risks of primary infection for different regions (Douma et al., 2016). Instead, we concentrate on the situation as faced directly before and after an initial incursion, restricting our attention to two representative 50km ***×*** 50km regions in Spain within which commercial citrus is grown at high-density and where HLB and/or AfCP and/or both could be introduced.

We demonstrate how an uncontrolled outbreak might spread if HLB entered, and how the speed of invasion would depend on whether AfCP was already locally widespread. We then investigate early detection surveillance, testing how the size of any epidemic at the point of first detection responds to the frequency and intensity of surveys. Finally, we assess the effectiveness of strategies for disease control, showing how relative efficacy can be quantified. By comparing results for two regions, we test the robustness of our conclusions.

## Methods

### Modelling spread of pathogen and vector

#### Citrus host distribution

Commercial and residential/municipal (henceforth “residential”) citrus were mapped across Spain and Portugal, and rasterised at 1km **×**1km resolution (Fig. 1(B); see also S1 Supporting Methods). For cell *i*, citrus densities (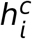 and 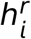 ) were converted into pairs of integer-valued numbers of “host units” (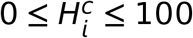 and 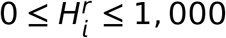) for compatibil-ity with our epidemiological model. We used different discretisations for commercial and residential citrus to allow our model to properly capture within-cell epidemiological dynamics, since within-cell densities are rather different (Fig. 1(C)), largely because commercial trees are typically planted in rows of hundreds/thousands of trees in close proximity. By separating commercial and residential citrus, our model can capture systematic differences in disease detection and control between settings.

**Figure 1:**
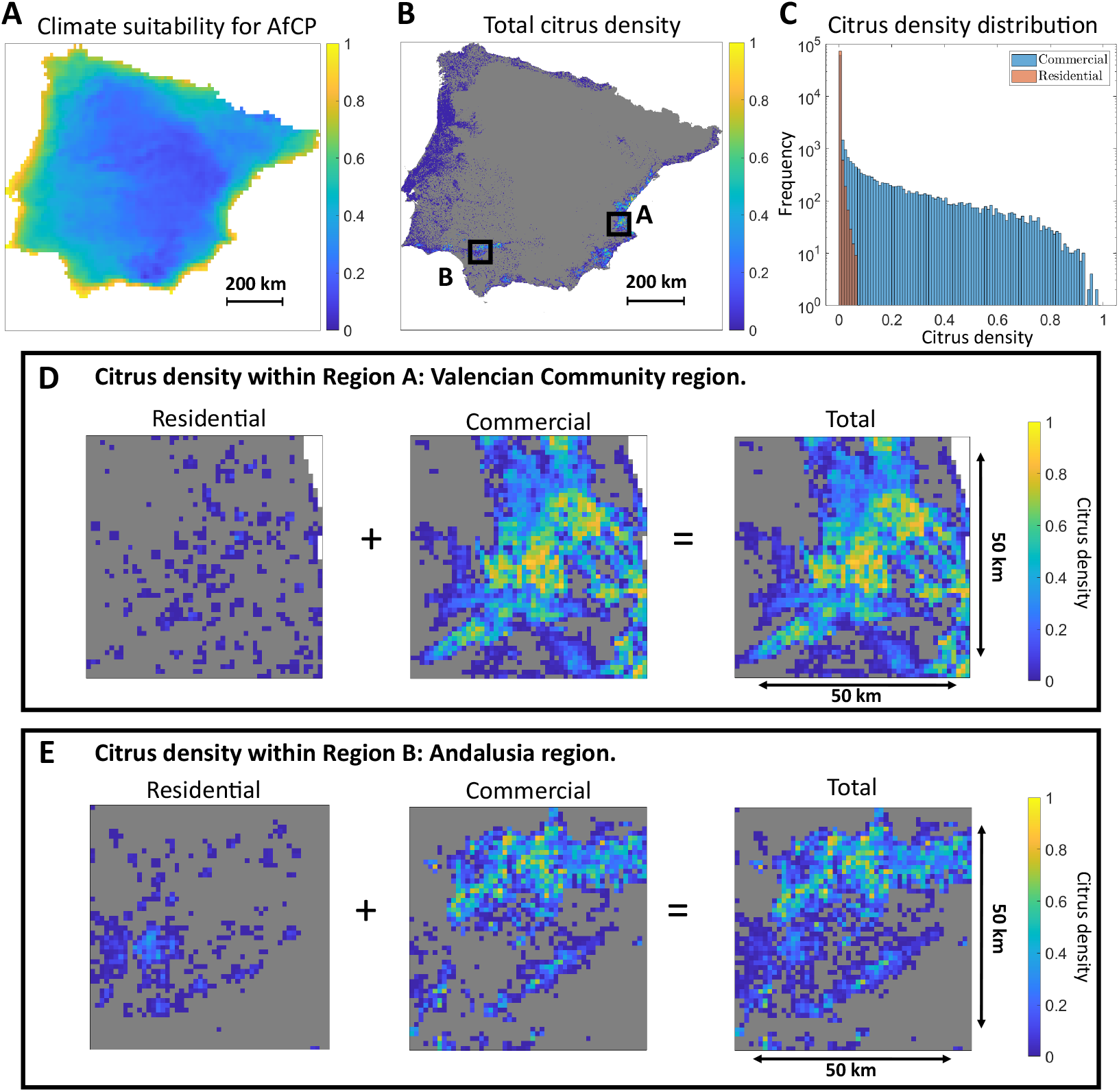
Climate suitability and citrus density across the Iberian peninsula. (A) AfCP climate suitability (see S1 Supporting Methods). (B) Total citrus density (residential **+** commercial) for each 1 km **×** 1 km cell, with our two 50 km **×** 50 km focal regions labelled. (C) Frequency distributions of (non-zero) residential and commercial citrus densities in each cell. (D) Residential, commercial and total citrus density maps (Region A), within the Valencian Community region, on the east coast of Spain. (E) Residential, commercial and total citrus density maps (Region B), within Andalusia, in southern Spain.

#### Bacterium and vector

We focus on the *C*Las bacterium as it is the most damaging, as well as the most likely to be introduced due to large and ongoing epidemics worldwide (Gottwald, 2010). We focus on AfCP as the vector, motivated by the availability of psyllid spread data from the recent invasion of northwestern Spain and Portugal (Cocuzza et al., 2017), and concomitant risk of spread of AfCP to regions of the Iberian Peninsula with commercial citriculture.

#### Disease and vector status within each cell

Our model tracks the HLB status of each citrus host unit in each cell, for both residential and commercial citrus. We distinguish: (*S*)usceptible, (*E*)xposed, (*C*)ryptic, (*I*)nfected and (*R*)emoved (Fig. 2). Susceptible host units are uninfected. Exposed host units are latently infected, i.e., not yet infectious. Cryptic host units are infectious but not yet symptomatic (Craig et al., 2018). Infected host units are infectious and symptomatic. Removed host units have been rogued (i.e., removed following detection to slow or stop the spread of disease). We do not account for other citrus demography, e.g., planting or disease-induced/natural death.

**Figure 2:**
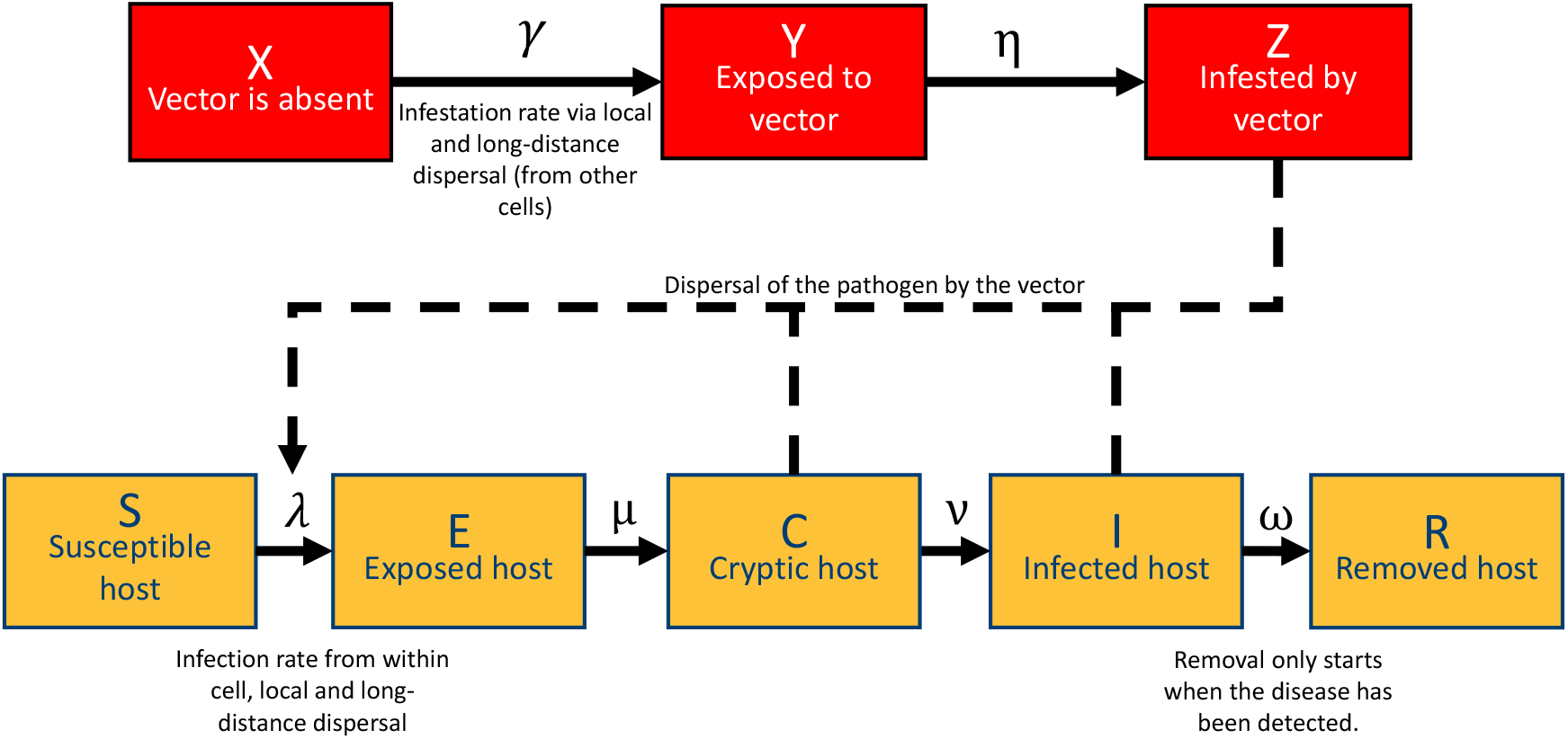
Model of vector (AfCP) and pathogen (*C*Las) in each 1km ×1km cell. We track each cell’s vector infestation status, for each class of citrus (residential/commercial), distinguishing: free of vector, *X*, exposed (psyllid is present, but has not yet fully colonised the cell), *Y*, and infested, *Z*. We also track the disease status of citrus within each cell, quantifying local densities of infection by tracking the number of host units in each epidemiological class in each cell, again distinguishing residential from commercial citrus. Epidemiological classes: susceptible (free from HLB), *S*; exposed (latently infected), *E*; cryptic (infectious but not symptomatic), *C*; infected (infectious and symptomatic), *I;* and removed, *R* (controlled by roguing). The rate from *X* **→** *Y, γ*, is the combination of local and long-distance dispersal of the vector (Γ, Eqns. 2-3 and Ω, Eqn. 4). The rate from *S* **→** *E, λ*, is the equivalent combination for infection (Λ, Eqns. 5-6, and Ω, Eqn. 4).

Epidemiological transitions of host units – and all other events in the Matlab implementation of our stochastic model – are simulated using Gillespie’s algorithm (Keeling and Rohani, 2008). Transitions from *E* **→** *C* and *C* **→** *I* occur at fixed rates *μ* and *ν*, respectively, with average latent period of 1 year, and average incubation period (i.e., to detectable symptoms) of 1.25 years (Parry et al., 2014) (Table 1). The rate at which host units transition from *S* **→** *E, λ*, is complex, and depends on psyllid dispersal and the infection status of citrus within the cell of interest and elsewhere, as described below. The transition from *I* **→** *R* occurs at rate *ω*, and depends on the detection and control strategy adopted, since it corresponds to roguing.

**Table 1:** Model parameters, descriptions, values and sources. Model fitting is described in overview in the main text, with further details in S1 Supporting Methods.

| Parameter |  | Estimate and source |  |
| --- | --- | --- | --- |
| $\delta$ | Psyllid dispersal rate | 1,600d<sup>-1</sup> | Fitted (see text) |
| $\zeta$ | Proportion of long-distance dispersal | 0.001 | Fitted (see text) |
| $\alpha_{loc}$ | Local dispersal scale | 1.96km | Nguyen et al. (2023) |
| $\alpha_{ld}$ | Long-distance dispersal scale | 130km | Benhadi-Marín et al. (2020) |
| $m$ | Vector reduction in commercial citrus | 0.9 | Qureshi et al. (2014) |
| $\eta$ | Rate of vector establishment | 1/365d<sup>-1</sup> | Assumed/fitted (see text) |
| $\beta$ | Infection rate | 10d<sup>-1</sup> | Fitted (see text) |
| $\rho$ | Proportion of within cell infection | 0.7 | Fitted (see text) |
| $\mu$ | Exposed to cryptic transition rate | 1/365d<sup>-1</sup> | Parry et al. (2014) |
| $\nu$ | Cryptic to symptomatic transition rate | 5/365d<sup>-1</sup> | Parry et al. (2014) |
| $\omega$ | Roguing rate | 0 | No control by default |

AfCP is currently only present in northwestern Spain and western Portugal, and so we model whether each cell’s citrus is infested by the psyllid. The model tracks whether cells are colonised by populations of psyllids that are sufficiently well-established for dispersal elsewhere, distinguishing infestation statuses of residential and commercial citrus. For each class of citrus, in each cell, there are three possibilities: vector absent (*X*), exposed (*Y*), and infested (*Z*). Exposed corresponds to a recently arrived vector population which is not yet capable of further dispersal. The rate of the *X* **→** *Y* transition, *γ*, is complex, and is described below. We assume the rate of the *Y* **→** *Z* transition is *η* **=** 1yr***−***1, corresponding to an average of one year for psyllid populations to fully colonise cells (see also S1 Supporting Methods).

Following colonisation, relative psyllid population densities in cell *i*, 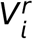and 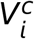, depend on local citrus densities and environmental condi-tions

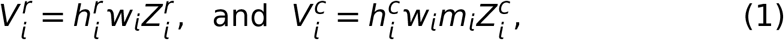

where 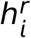 and 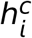 are proportions of residential/commercial citrus 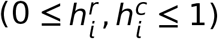 and *W*_*i*_ is the climate suitability for psyllids (0 ***≤*** *W*_*i*_ ***≤*** 1; see also S1 Supporting Methods). For commercial citrus, the effect of pest management, *m*_*i*_ is also included (0 ***≤*** *m*_*i*_ ***≤*** 1; see also S1 Supporting Methods, and note our mapping procedure allows us to account for a lack of management in abandoned and/or organic orchards). In Eqn. 1 the infestation status (i.e., 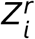 or 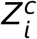) acts as an indicator function to ensure psyllid densities are only non-zero when the cell is fully colonised.

#### Interactions between cells

##### Scales of dispersal

We distinguish two scales of dispersal. The local dispersal kernel, 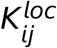, reflects psyllid movement between an individual cell, *i*, and one of its near neighbours, *j*. For this we use an exponential kernel as fitted to spread of ACP in the United States by Nguyen et al. (2023). However, psyllids can occasionally travel much further, due to extreme wind (Antolinez et al., 2021) or human transportation (Nunes et al., 2023). We capture these relatively infrequent long-distance dispersals using *t*-distribution kernel as fitted to the AfCP invasion in Portugal by Benhadi-Marín et al. (2020). Mathematical details are in S1 Supporting Methods.

##### Dispersal of psyllids

Local dispersal of psyllids (AfCP) occurs from populations which have colonised neighbouring cells, with local forces of infestation on cell *i*

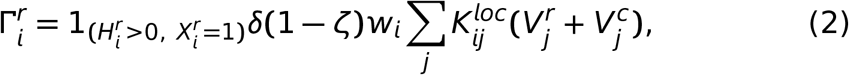

and

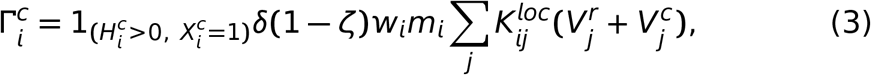

where 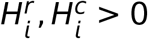 and 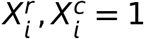 ensure only cells containing citrus but currently psyllid-free can become infested, 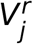 and 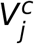 are densities of (established) psyllid populations in residential/commercial citrus in cell *j*, 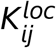 is the local dispersal kernel, *δ* is the rate of psyllid dispersal, and *ζ* is the proportion of long-distance dispersals. For both residential and commercial citrus, forces of infestation include *W*_*i*_, representing climatic effects. For commercial citrus, *m*_*i*_ is also included in Eqn. (3), representing reduced establishment probability due to pest management.

Our model implements the “particle-emission” formulation of long-distance dispersal (Meentemeyer et al., 2011). Rates of long-distance dispersal from cell *i* are

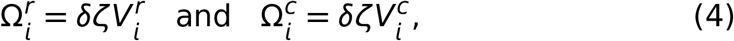

where *δ* is the dispersal rate, *ζ* is the proportion of long-distance dispersals, and 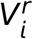 and 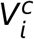 are densities of psyllid populations in residential/commercial citrus in cell *i*. Whenever a long-distance dispersal event is sampled by our Gillespie algorithm, the angle of dispersal is drawn uniformly on [ 0, 2*π*), and a distance sampled from the long-distance dispersal kernel. If the corresponding destination cell, *j*, contains citrus, the type of citrus challenged (residential or commercial) is randomly chosen according to the proportion of each type. The probability the vector will infest is given by *W*_*j*_ for residential citrus, or *W*_*j*_*m*_*j*_ for commercial citrus. If cell *j* does not contain citrus, the vector simply fails to disperse.

##### Spread of infection

Infection rates are closely coupled to psyllid dispersal. However, since our model captures HLB dynamics within individual cells, within- and between-cell infection must be distinguished. The rate of local infection of susceptible host units in residential citrus in cell *i* (i.e., the component of the rate of the *S* → *E* transition in cell *i* corresponding to within-cell and nearby sources of infection) is

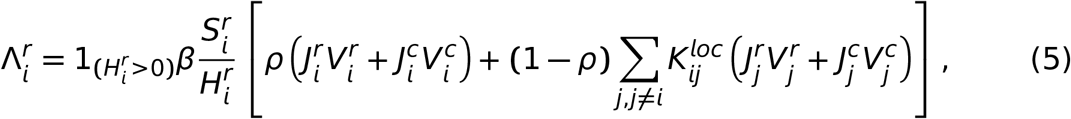

and for commercial host units is

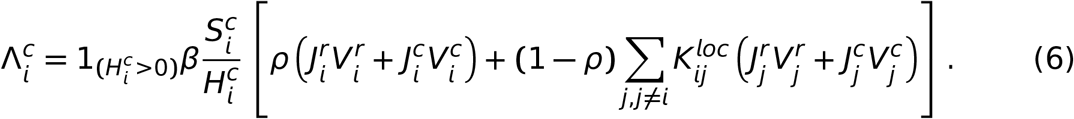

In Eqns. 5 and 6, *β* is the infection rate, 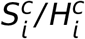 and 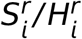 are propor-tions of uninfected commercial/residential host units in cell *i, ρ* is the proportion of within-cell transmission, and 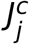 and 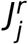 are proportions of infectious commercial/residential citrus in cell *j*, i.e.,

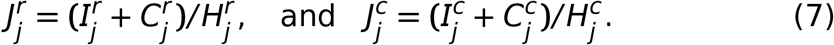

Long-distance transmission of HLB is a consequence of psyllid movement. Whenever long-distance psyllid dispersal occurs from residential citrus in cell *i*, the probability the recipient cell (*j*) gains a single HLB exposed host unit is given by 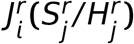 or 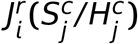, depend-ing on whether the long-distance dispersal challenges residential or commercial citrus. Analogous probabilities involving 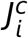 set the chance of infection from long-distance dispersal from cell *i* originating within commercial citrus. There is no requirement for a vector population to establish in the recipient location for transmission of the bacterium, and so parameters for climate and pest management are not included in these probabilities. However, of course, infection will not be able to spread onwards from any infected cell until a psyllid population does colonise locally.

##### Parameterisation

We fix values of five parameters in our model from the literature (Table 1): *α*_*loc*_ (scale of local psyllid dispersal); *α*_*ld*_ (scale of long-distance psyllid dispersal); *m* (effect of insecticide sprays on psyllid populations); *μ* (rate at which exposed hosts become cryptic); and *ν* (rate at which cryptic hosts become symptomatic). The rate of roguing, *ω*, depends on the control strategy, and in model runs without disease control we assume *ω* **=** 0.

However, five remaining parameters are fitted to data (S1 Supporting Methods). These are: *δ* (dispersal rate of psyllids); *ζ* (proportion of long-distance psyllid dispersal); *η* (rate at which psyllids fully colonise a cell following first infestation); *β* (pathogen transmission rate); and *ρ* (rate of within-cell relative to between-cell infection). Parameters *δ* and *ζ* are fitted to AfCP presence data from surveys in Portugal and Spain up to 2021 (Benhadi-Marín et al., 2022), contingent on an assumed value of *η*. To fit parameters *β* and *ρ*, we calibrated results our model against a previous model of the spread of HLB (*C*Las vectored by ACP) in Florida (Mastin et al., 2020). Full details are in S1 Supporting Methods.

### Modelling detection and control

#### Early detection

Before first detection, we model regular inspections every Δ (“inspection interval”) years, with *a*^*c*^% and *a*^*r*^ % of cells containing commercial and residential citrus, respectively, across our 50km **×** 50km region, randomly selected on each round of inspection. Selection is weighted by within-cell citrus density. Within each selected cell, at any inspection, *n*_*h*_ host units are selected at random (if the cell has fewer than *n*_*h*_ host units of the prescribed type, all are inspected). Disease is detected with probability *p* on each symptomatic (i.e., class *I*) host unit. The first inspection is at a random time on [ 0, Δ), where 0 is the time of first introduction.

#### Control

Following first detection, we assume inspection significantly intensifies, occurring according to the roguing interval, Δ_*R*_ (generally with Δ_*R*_ *<* Δ). This detection regime applies across the region, and so we assume the entire 50km **×** 50km area corresponds to the Infested Zone under Regulation (EU) 2016/2031 (European Union, 2016). We assume the increased threat of disease encourages most stakeholders to participate in enhanced detection and control. However, since some growers will not cooperate, we introduce a compliance parameter, *c*, and assume only a proportion *c* of growers comply. The set of non-complying growers remains fixed for each simulation. We assume that detection and control does not occur within the cells containing non-compliant growers. However, we assume all host units in all complying cells are inspected on each roguing interval.

Detection of symptomatic host unit(s) triggers roguing (i.e., removal of host). Following detection, which as for early detection occurs with probability *p* for symptomatic host units, each detected host unit is (immediately) removed with probability *q*. With probability 1 **−** *q* a host unit will not be removed, although of course it may be re-detected and removed later. The roguing probability, *q*, therefore acts as a proxy for any delays in control and/or imperfect management by growers or plant health authorities. We also allow for commercial growers applying extra pest management following first detection. This reduces the vector population by a further factor *m*\*, over-and-above the reduction by *m* caused by “standard” pest management. We model this by assuming *m* is increased by (1**−** *m*)*m*\* across the entire region of interest immediately after first detection, and that this decreases vector populations in commercial citrus (while accounting for abandoned/organic citrus, and only for the subset of growers who comply).

#### Parameterisation

Since *C*Las is an EU priority pest (European Union, 2016, 2019), annual surveys are mandatory, and so inspection is assumed once per year (Δ **=** 1yr). We assume by default (see also Table 2) *a*^*c*^ **=** 1% of cells with commercial citrus are (randomly) surveyed each year, and within each cell *n*_*h*_ **=** 5 (commercial) host units are inspected. However, by default residential citrus is not inspected (*a*^*r*^ **=** 0%), reflecting the difficulty of surveillance in private gardens and other residential settings (Cocuzza et al., 2017). We assume the probability of (visual) detection of symptomatic hosts is *p* **=** 0.5 (Mastin et al., 2020).

**Table 2:**
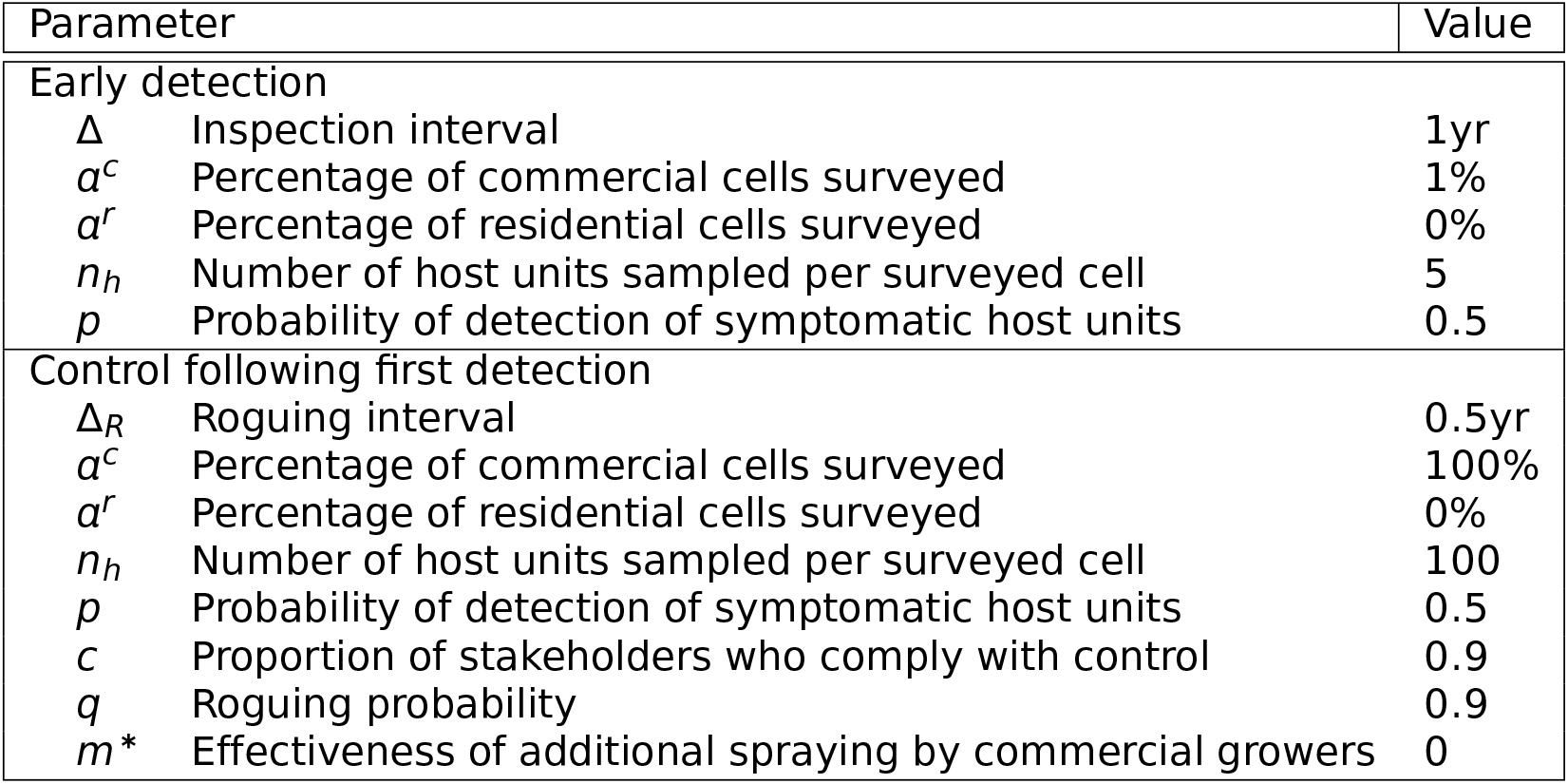
Default parameters for detection and control strategies. There is initially an early detection phase, but following detection anywhere across the region, the strategy switches to include more intensive surveillance and active disease control (details in text).

After detection of the pathogen anywhere within the region, our baseline is that the default compliance rate of commercial growers is 90% (i.e., *c* **=** 0.9). For those growers who comply, the roguing interval is Δ_*R*_ **=** 0.5 (i.e., detection/control every 6 months) and all commercial citrus units are inspected within all cells (*n*_*h*_ **=** 100, *a*^*c*^ **=** 100%). The roguing probability is *q* **=** 0.9. However, we assume residential citrus remains uninspected 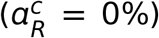; because of this, by default we assume no roguing is done for residential citrus. We also assume there is no additional pest management introduced by commercial growers (i.e., *m*^*^ **=** 0). However, our sensitivity analyses allow consequences of these choices to be tested.

## Results

### Disease spread without control

We initially focus on Region A (Fig. 1(D)), in the Valencian Community region (eastern Spain), one of the main citrus growing regions in the EU. If both the vector and HLB are introduced simultaneously into a single cell, HLB spreads rapidly (Fig. 3 and S3 Supporting Videos (Video 2)), although spread of the vector is – of course – even faster (S2 Supporting Results, Fig. S10, and S3 Supporting Videos, Video 1). Several cells near to the point of initial introduction and two cells which are further away become HLB positive within the first year in this single simulation (Fig. 3(A)). Almost all cells with citrus have at least one infected citrus host unit by year 10 (Fig. 3(D)), and within 20 years almost all susceptible citrus across the entire region is infected (compare Fig. 3(E) with Fig. 1(D)).

**Figure 3:**
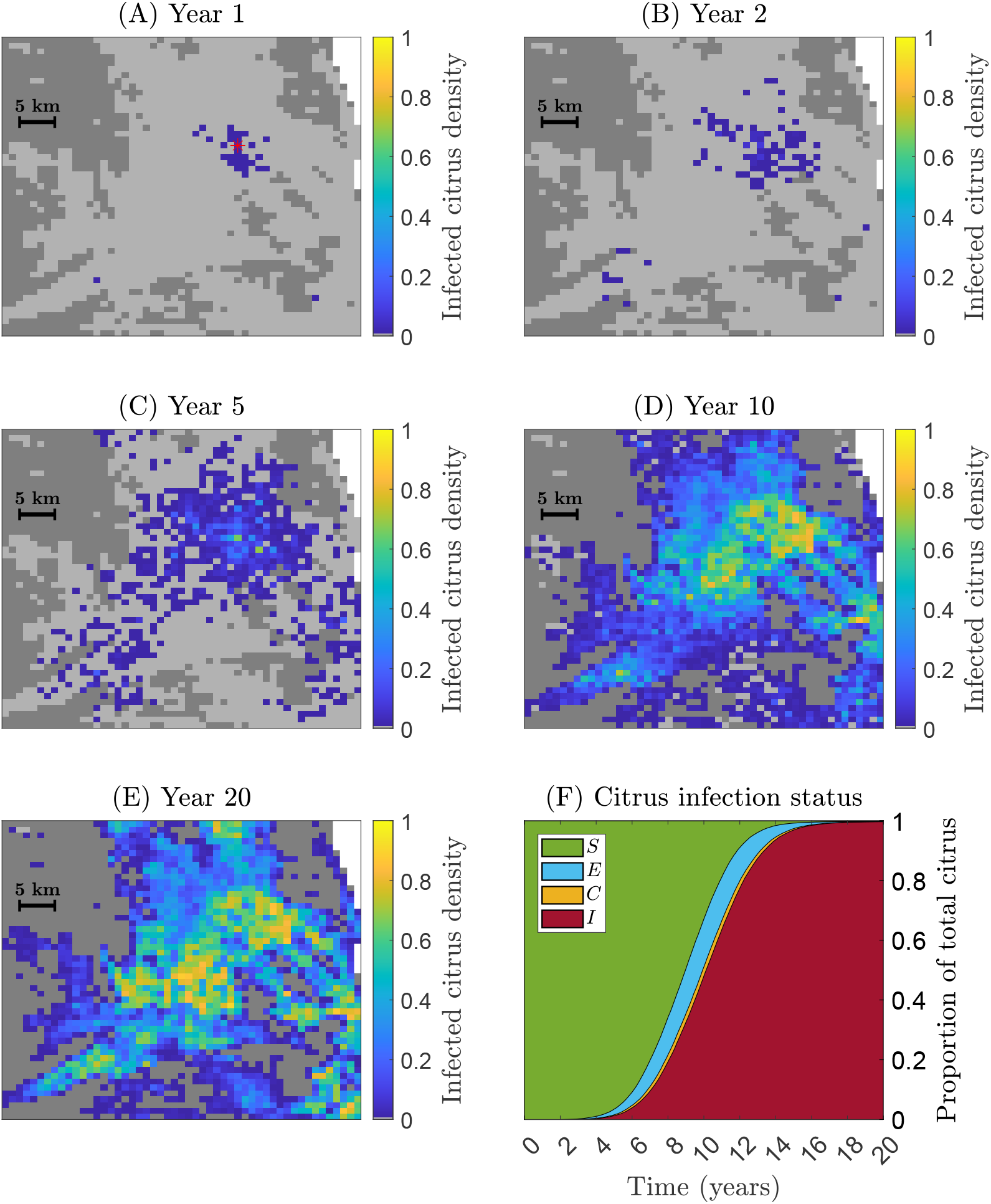
Spread of the pathogen in a single simulation in Region A (Valencia). Both vector and pathogen are introduced simultaneously at *t* **=** 0 into a single 1km **×** 1km cell (asterisked in panel (A)). A single host unit of commercial citrus is moved from *S* **→** *E*, and the psyllid status of commercial citrus in that cell set to *Y*. (A)-(E) Maps showing the density of infected citrus host units (*E* **+** *C* **+** *I*) within each cell at different times after introduction; see also S3 Supporting Videos (Video 2). (F) Disease progress curve showing proportions of citrus over the entire region in each epidemiological compartment, (S)usceptible, (E)xposed, (C)ryptic and (I)nfected (no host units are (R)emoved, since disease control by roguing was not done in the underpinning simulation).

Although individual simulations are easy to visualise, only ensembles of multiple simulations capture the range of possibilities from our stochastic model. Although the vector spreads rapidly, and it does not take long for an initial vector population to become widely dispersed, if the vector is already widespread at the time of pathogen entry, HLB invasion is faster, since spread can begin immediately (compare Figs. 4(A) and 4(B)). Prediction intervals are wider when the vector must also spread, since the variability in the spread of the vector has a knock-on effect upon HLB. This is particularly pronounced when the vector is (randomly) introduced into a cell with low density citrus.

**Figure 4:**
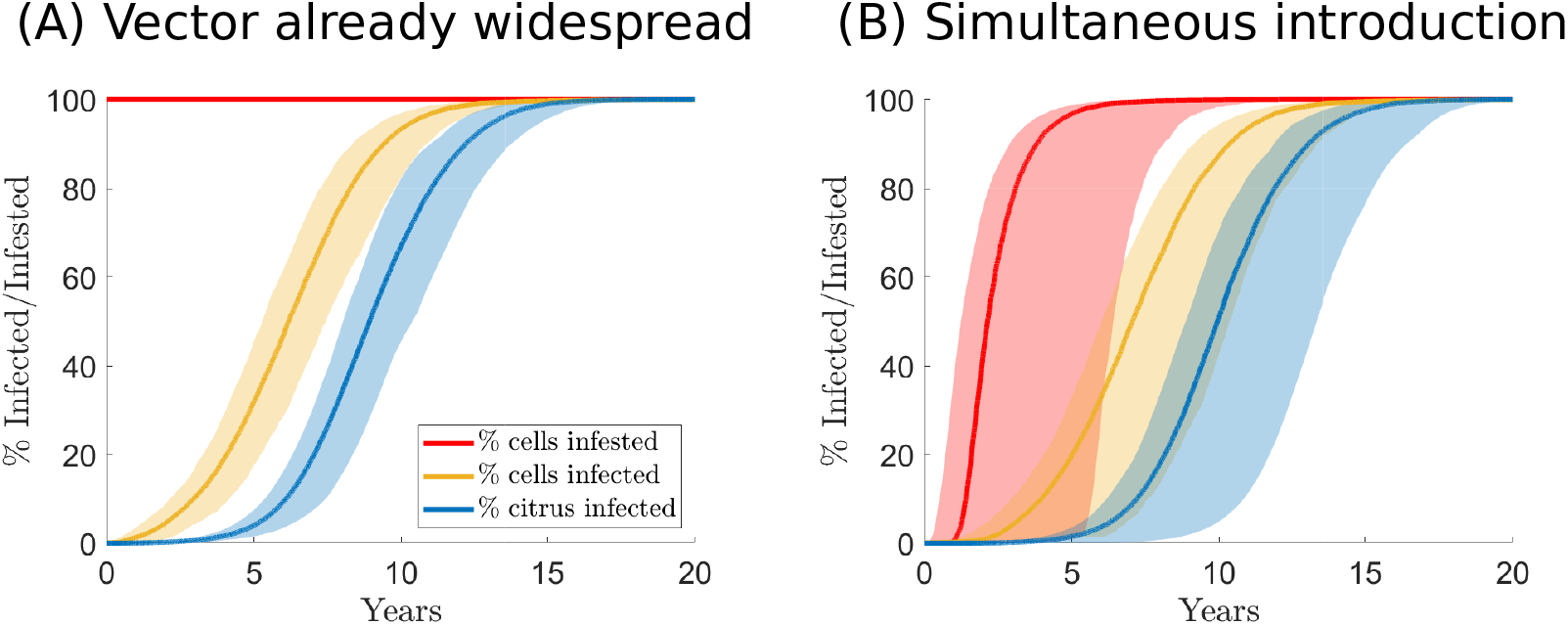
Effect of prior invasion by the vector on pathogen spread in Region A (Valencia). The percentage of cells infested by the vector (red), with at least one infected unit of citrus (yellow) and the percentage of all citrus infected (blue), when the vector is already widespread throughout the region of interest (Panel (A)) *versus* when the vector and pathogen are introduced simultaneously (Panel (B)). In all cases the pathogen is introduced into the same cell at *t* **=** 0; for the simulations shown in Panel (B), the cell initially infested by the vector is chosen at random from all cells containing commercial citrus. Shaded regions show 95% prediction intervals from an ensemble of 200 simulation results.

Henceforth, we focus exclusively on the case in which the vector is already widespread in the region of interest. However, in all subsequent simulations, the initial location of *C*Las infection is selected at random (weighted by commercial citrus density, assuming that the pathogen is introduced on planting material, and that higher density commercial operations plant larger amounts more frequently).

### Early detection surveillance

We test early detection surveillance by varying the proportion of commercial cells inspected on each survey (*a^c^*), and the interval between successive surveys (Δ), considering three indicative probabilities of detection: *p* **=** 0.2, 0.5 and 0.8. We vary *a^c^* from 0.5% (9 cells across Region A), to 5% (83 cells). We vary the inspection interval (Δ) from once every 3 months to once every 2 years. The number of host units of citrus to inspect per cell is fixed at the default value *n*_*h*_ **=** 5 (this is relaxed in S2 Supporting Results, Fig. S11).

We summarise performance via the time until first detection and proportion of citrus then infected (Fig. 5). Unsurprisingly, effective early detection is conditioned on inspecting as many cells as possible, as often as possible (Figs. 5(A) and (D)). However, we note the variability in both time of first detection (Figs. 5(B) and 5(E)) and (particularly) the proportion of citrus infected (Figs. 5(C) and 5(F)) is larger when the probability of detection, *p* is lower. There are limited returns from increasing the probability of detection if that probability is already relatively high (compare larger differences between responses for *p* **=** 0.2 and *p* **=** 0.5 to smaller differences between those for *p* **=** 0.5 and *p* **=** 0.8 in Figs. 5(B), 5(C), 5(E) and 5(F)).

**Figure 5:**
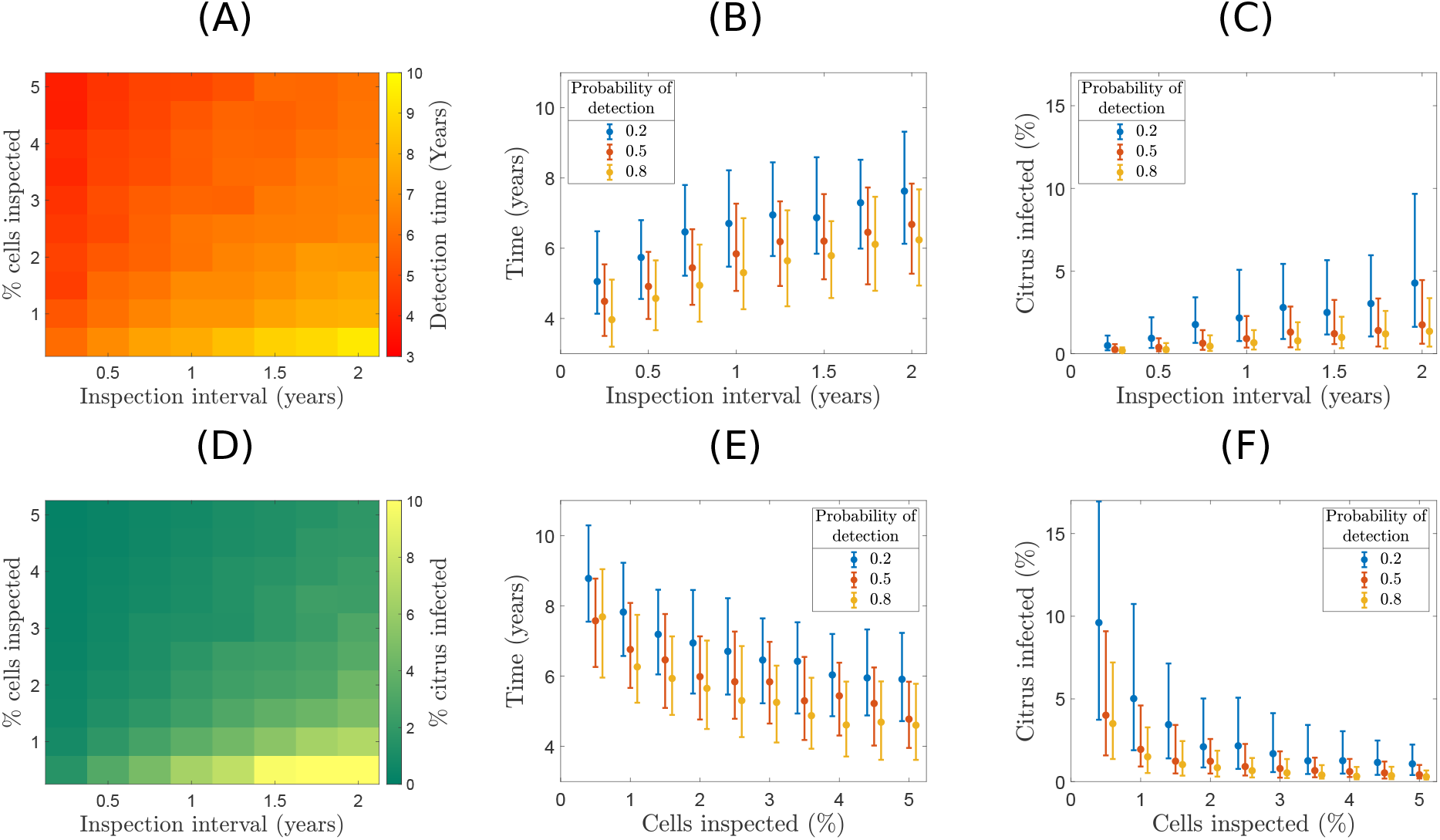
Effectiveness of surveillance strategies in Region A (Valencia). (A) Mean time of detection and (D) Mean percentage of total citrus infected at the time of detection. The number of host units sampled in each surveyed cell is *n*_*h*_ **=** 5 and the probability of detection of symptomatic host units is *p* **=** 0.5. (B,C,E,F) Median and inter-quartile ranges of (B,E) time until detection and (C,F) proportion of citrus that is infected. The inspection interval and percentage of cells inspected are varied; (B,C) has fixed 5% of cells inspected at varying intervals, and (E,F) varies the percentage of cells inspected at a fixed 12 month inspection interval. Plots show responses for three values of the probability of detection, *p* **=** 0.2, 0.5 and 0.8. Results are shown from ensembles of 200 simulations for each set of parameters.

### Effectiveness of control

#### Default parameterisation

We examine first a single simulation using the baseline strategy (Table 2), in which early detection surveillance is followed by more intensive detection and control once the pathogen is detected (Fig. 6 and S3 Supporting Videos (Video 3)), noting that results in this single simulation are typical of those from a much larger ensemble (see also S2 Supporting Results, Fig. S12). Although roguing – which in this particular simulation started following detection around 5 years after first introduction – slows pathogen spread (compare Figs. 3 and 6), after 20 years 65% of all citrus host units have been removed, a further 12% of units are infected, and the epidemic is still ongoing, although spreading relatively slowly. Cells containing high densities of infected citrus (green/yellow cells in Fig. 6(E)) are those within which commercial growers are non-compliant, and so from which no infected citrus is removed. Such uncontrolled locations act as a source of inoculum driving the ongoing epidemic.

**Figure 6:**
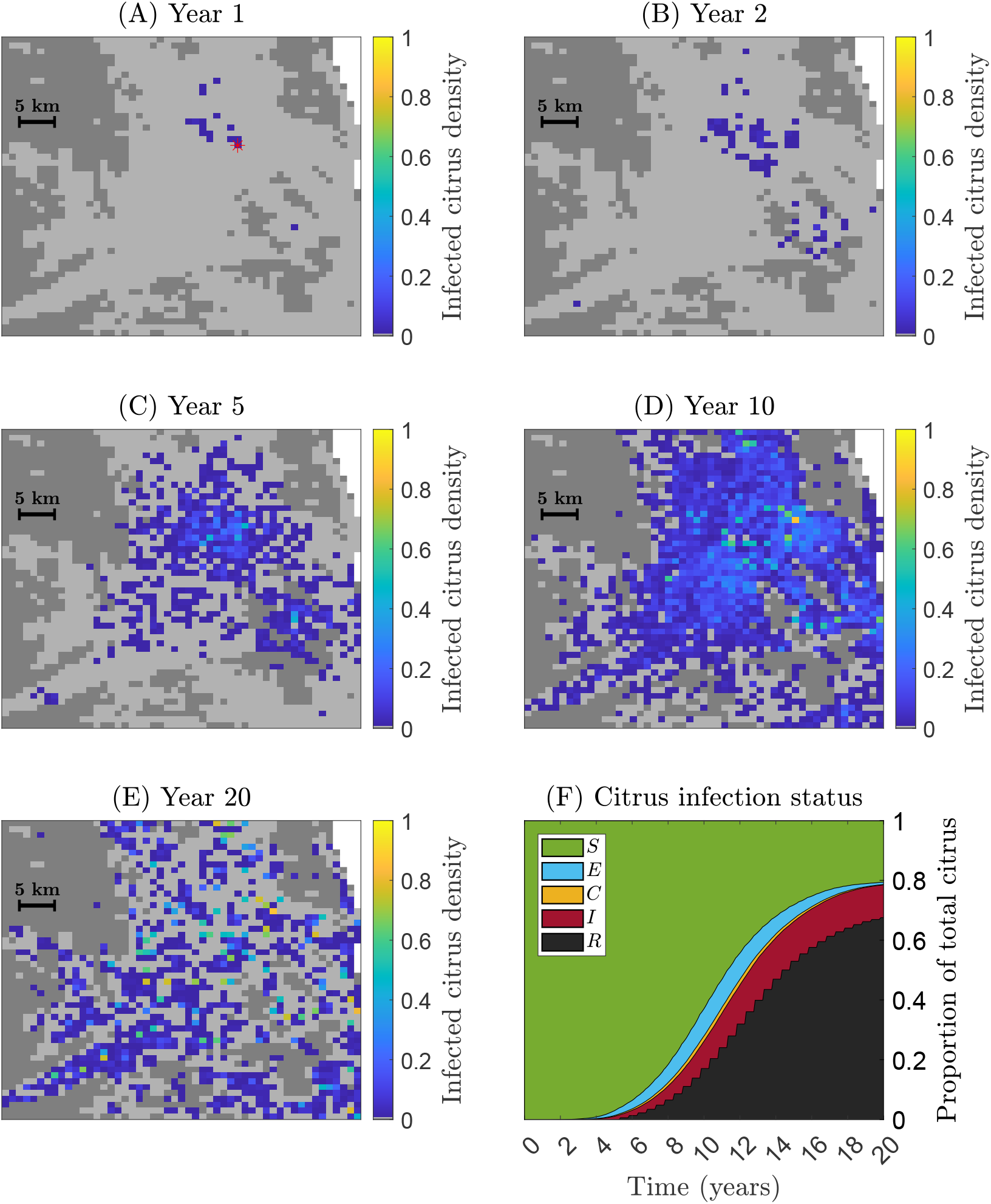
Spread of the pathogen in a single simulation in Region A (Valencia) using baseline parameters for detection and control (Table 2). Initial conditions as Fig. 3. In this simulation, disease is first detected after approximately 5 years, after which control started immediately. Maps show densities of infected citrus (*E* **+** *C* **+** *I*) in each 1km **×** 1km cell; see also S3 Supporting Videos (Video 3).

#### Sensitivity analysis

We do a series of one-way sensitivity scans, examining percentages of infected/removed citrus over time when varying single parameters in ensembles of simulations (Fig. 7). These scans conveniently summarise the relative importance of different aspects of disease management. Reducing the roguing interval (Δ_*R*_) and the compliance and roguing parameters (*c* and *q*) have strong effects on the epidemic (Figs. 7(A), (B) and (E)), as these parameters directly impact the rate and/or number of units of citrus removed. Conversely, the proportion of cells inspected in the early detection strategy (*a^c^*) does not have a substantial effect on epidemic rates (Fig. 7(F)), since it only affects the time of first detection. Roguing commercial citrus is essential (Fig. 7(C)), since the vast majority of citrus is commercial, although additional management of residential/municipal citrus leads to a visible difference in epidemic progression.

**Figure 7:**
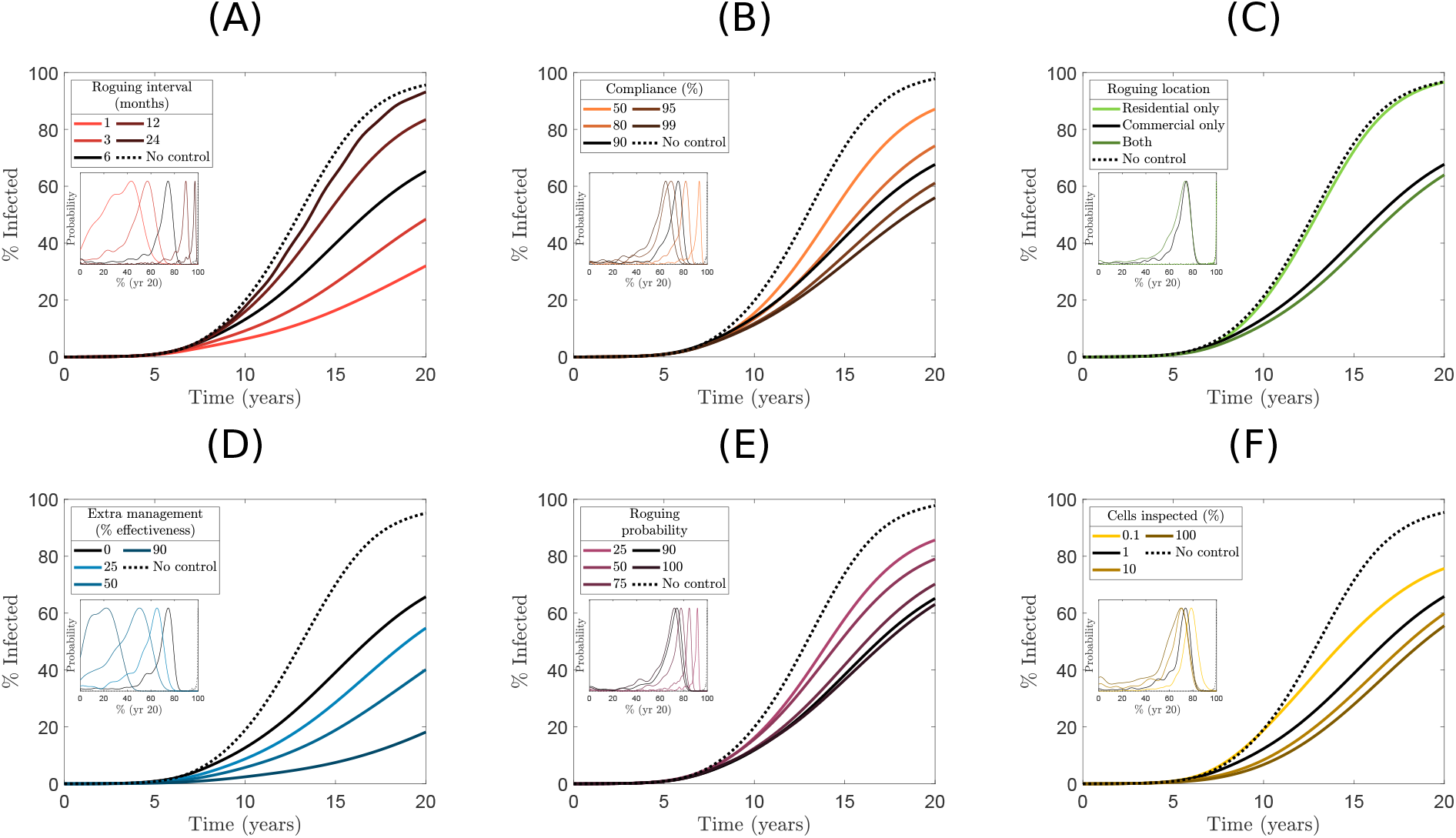
Management scenarios in Region A (Valencia). Mean proportions of infected or removed citrus (*E* **+** *C* **+** *I* **+** *R*) over time when varying a single parameter from the baseline parameterisation (Fig. 6 and Table 2). Aspects varied in each panel: (A) roguing interval, (B) grower compliance, (C) types of citrus rogued, (D) increases to the pest management parameter, *m*^*^, (E) roguing probability, and (F) proportion of cells inspected for early detection. Inset graphs show probability distributions of the proportion of infected or removed citrus after 20 years (normalised to have the same maximum for ease of visualisation). Black lines show results using the baseline parameterisation; dotted lines show results with no control. Averages and probability distributions calculated from ensembles 200 simulations per parameter combination.

However, the parameter with the strongest impact on control effectiveness is *m*\*, the effectiveness of additional pest management to reduce the size of the vector population once the pathogen is detected (Fig. 7(D)). Even before detection, we assume routine insecticide sprays lead to a *m* **=** 90% reduction to vector densities in commercial citrus. However, if HLB detection triggers additional pest management, and if the psyllid population is reduced by up to 99%, this can slow the progression of the epidemic, although it does not stop spread completely. We return to the plausibility of such high levels of vector control in the EU below.

#### Robustness

We repeat our analysis of control efficacy for Region B (Fig. 1(E)), in the Andalusia region (southwestern Spain). Region B has less high-density commercial citrus than Region A, although still contains substantial production, and more residential citrus that may hinder attempts to control spread. Region B is also much closer to where AfCP has already been found, and so may be at higher risk. However since Region B is further inland, there is a lower climate suitability compared to Region A. Fig. 8 shows the importance of each parameter, equivalent to Fig. 7 for Region A (further results for Region B are in S2 Supporting Results, Figs. S14-S18, and S3 Supporting Videos, Videos 4-6). Impacts of control measures are remarkably similar between the two regions, and additional pest management (*m*\*) remains the most effective intervention.

**Figure 8:**
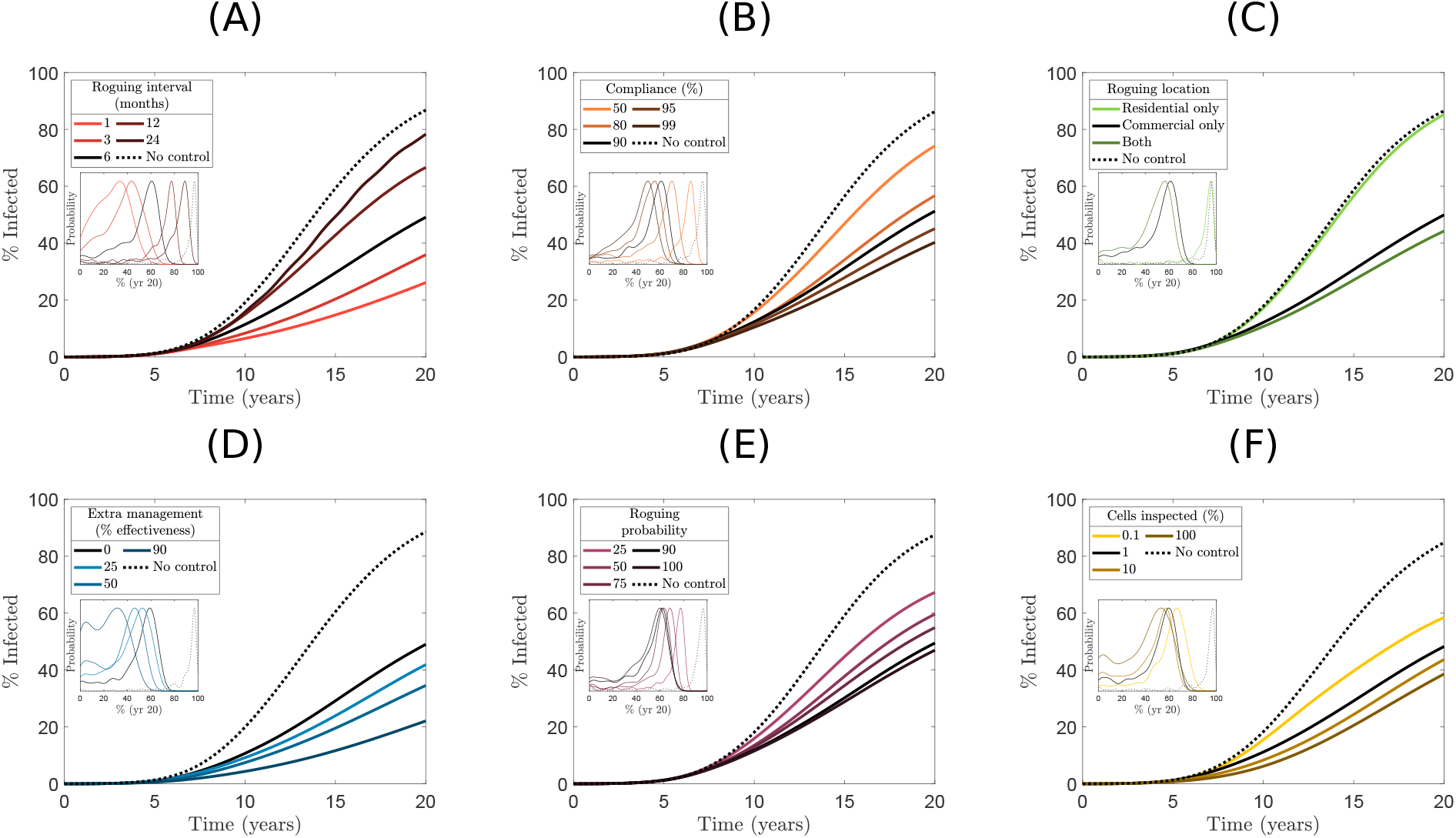
Management scenarios in Region B (Andalusia). Mean proportions of infected or removed citrus (*E* **+** *C* **+** *I* **+** *R*) over time when varying a single parameter from the baseline case (Table 2). Individual panels as Fig. 7.

## Discussion

HLB has not been reported in the EU, although invasion is possible, perhaps even probable (Wang, 2020). We demonstrate how mathematical modelling can contribute to developing epidemiological preparedness for a future HLB invasion, focusing on regions in Spain containing high-density citrus production. We found that any epidemic is likely to be very well-established at the time of first detection. Even with intensive disease management with almost all commercial growers participating in a large-scale programme of detection and roguing (i.e., removal of infected citrus trees), eradication is very likely to be impossible. However, a combination of sustained and rapid disease control via roguing and very heavy insecticide sprays may provide relatively good control over sustained periods, allowing citrus production to be maintained for some time. This echoes the experience of growers and regulators at least some other countries, most notably Brazil (Bassanezi et al., 2020).

Significant uncertainty surrounds the invading vector and bacterium. Although our model is transferable to different vector-bacterium combinations, we focused on invasion by *C*Las vectored by AfCP. Of the three *C*L species, *C*Las is the most widely distributed and damaging (Gottwald, 2010). Our choice should therefore be uncontentious. For the vector, we focused on AfCP, motivated by its presence in Portugal and Spain, and despite recent reports of ACP from Israel (EPPO, 2022) and Cyprus (EPPO, 2023). Focusing on AfCP allowed us to use data from Spain and Portugal to parameterise long-range psyllid dispersal in our model. Although both vectors transmit *C*Las (Reynaud et al., 2022), systematic differences in *C*Las infection rates are not well-characterised. However, since our model explicitly includes vector climate suitability (Fig. 1(A)), we account for AfCP’s relative lack of heat tolerance (Paiva et al., 2020). We used parameters and outputs from models of *C*Las vectored by ACP (Mastin et al., 2020; Nguyen et al., 2023) to parameterise short-range AfCP dispersal and HLB transmis-sion rates. While this was unavoidable, given the paucity of other information, it potentially understates biological differences between vectors.

Uncertainty also surrounds where HLB will first enter the EU. Although risks of entry have been modelled for the USA (Gottwald et al., 2019), there are no equivalent models for the EU. We therefore chose to focus on a reasonable worst-case scenario, with HLB introduced into high-density commercial citrus in Spain, the largest EU producer (Schimmenti et al., 2013). We compared results for two 50km **×** 50km regions, in Valencia and Andalusia (Fig. 1), and (generally) assumed AfCP was already locally widespread at the time of HLB invasion. Our approach was intended to put limits on the potential efficacy of control. If the vector were also actively spreading in whichever area HLB was invading, as is arguably more likely, any outbreak would proceed slightly more slowly (Fig. 4). However, relative efficacies of different management strategies are unaffected by prior invasion of the vector (S2 Supporting Results, Fig. S13).

In fact, while AfCP has spread widely in coastal Portugal and north-western Spain in the decade since detection (Perez-Otero et al., 2015; Siverio et al., 2017; Benhadi-Marín et al., 2022), it has not reached the main commercial citrus areas. Furthermore, (classical) biological control via the parasitoid *Tamarixia dryi* has slowed or even stopped spread in residential settings since 2019 (Molina et al., 2021; Duarte et al., 2024), although how *T. dryi* would be affected by insecticide sprays in commercial citrus remains unclear. Simultaneous invasion of vector and pathogen has been common previously. For example, in California, ACP was first detected in 2008 and HLB in 2012 (Nguyen et al., 2023), while in Florida, ACP was detected in 1998 and HLB in 2005. However, the pathogen is harder to detect than the psyllid, and there is consensus HLB was widespread in Florida by 2005 (Halbert et al., 2010).

The pathogen spreads rapidly in our model following first introduction (Fig. 3). We note a recent expert knowledge elicitation exer-cise (EFSA et al., 2019a), which estimated a median spread rate of 20.61km yr***−***1 (1**−** 99% range 0.90 **−** 40.12km yr***−***1) for HLB in the EU. While our spread rates are within this range, they lie towards the lower end, particularly early in invasions. There is a lag phase of a few years in which spread is relatively slow (Fig. 4), particularly when the pathogen is introduced into cells with low citrus density. Of course, too, we might also note sustained spread at 20km yr***−***1 would be impossible to discern at the scale we have focused on here. By restricting our attention to 50km **×** 50km regions, we have tended to de-emphasise effects of long-distance dispersal, even though this is included in our model, and at larger scales this would permit the pathogen to spread even more rapidly than 40km yr***−***1. Given rates of long-distance psyllid spread in our model were calibrated to be sufficient to replicate the invasion over hundreds of kilometres of coastal Portugal and northwestern Spain within only 6 years, it is important to note that impacts of HLB invasion would rapidly be realised far outside the 50km **×** 50km regions of initial invasion we focused on here.

The delay before first detection of HLB depends on surveillance intensity, but is 3 **−** 10 years for all parameterisations tested (Fig. 5). The range is similar to, but the average again slightly lower than, estimates reported following the expert knowledge elicitation exercise (EFSA et al., 2019a), i.e., a median of 2.1 years (1 **−** 99% range 0.6 **−** 6.7 years). The lower bound of 3 years for detection in our model largely reflects the short lag before rapid spread in our model, as discussed above. We assumed relatively large proportions of citrus were regularly being surveyed, and scaling this from our 50km **×** 50km regions to entire countries would be expensive. However, given we had no risk of entry based reason other than high-density commercial citrus to focus on the particular regions considered here, detecting the pathogen within these timescales would require an equally intensive country-wide survey, at least in regions of high citrus density. However, since all three *C*Ls are EU Priority Pests, annual surveys are required in every member state (European Union, 2016, 2019), and we note significant surveillance is currently mandated in the USA and Brazil (Parnell et al., 2014; Bassanezi et al., 2020). In our model, cells are chosen for inspection at random, weighted by citrus density. However, other strategies, such as targeting locations at higher risk, may detect the disease earlier or with less cost (Mastin et al., 2020; Parnell et al., 2014). Testing this would be most informative if done over larger spatial scales, driven by a model quantifying relative entry risks (Douma et al., 2016).

We modelled an immediate shift of strategy following detection, increasing surveillance and introducing disease control region-wide. The entire 50km **×** 50km region was therefore treated as the Infested Zone under Regulation (EU) 2016/2031 (European Union, 2016). This is a simplification of current plans, which are based on bounding the infected area via a delimiting survey. Implementation of such surveys has become increasingly statistical (EFSA et al., 2020), and is now based on identifying sample sizes required for a certain confidence in detection given an assumed disease prevalence. Recent work has tested performance of strategies for *Xylella fastidiosa* using a (small-scale) individual based model (Cendoya et al., 2024). Doing this for HLB would be interesting, and suitable small-scale models are already available (e.g., Parry et al. (2014); Craig et al. (2018)), although recalibration would be needed for use in the EU.

Roguing does not stop the epidemic but slows spread (Fig. 6). One driver is asymptomatic infection, since hosts are only removed once symptoms are detectable. Another is that we assume growers do not always comply with control, since HLB infected trees continue to produce fruit, at least for a few years (Bassanezi et al., 2011). Furthermore, private gardens, backyard trees and abandoned orchards act as refugia (Cocuzza et al., 2017). During an outbreak there will be several locations where management does not occur, and these fuel spread. Nevertheless, even with perfect compliance by commercial growers and active management of residential citrus, transmission is likely to continue (Figs. 7 and 8). This is due to the cryptic pe-riod within which plants are infectious but not symptomatic, and so not detected/removed. Based on previous model fitting (Parry et al., 2014), we used a relatively lengthy asymptomatic period (1.25 years on average; Table 1). Detecting HLB before visual symptoms would improve performance, even if diagnostic tests were inefficient (Mastin et al., 2022). Impressive results have been reported from Florida using dogs trained to identify infections before symptoms are visible (Gottwald et al., 2020). However, transferability and application over large spatial scales remain to be tested.

Asymptomatic infection means host removal could also be improved by removal of all trees within a particular radius of detected infection (Cunniffe et al., 2015b). This is implicitly accounted for via our host quantisation, since roguing removes entire commercial host units (i.e., areas of 100m **×** 100m = 1 ha). However, modelling different radii of removal is relatively simple (Cunniffe et al., 2016; Hyatt-Twynam et al., 2017), and would be an interesting extension, perhaps particularly if coupled to more detailed models of grower behaviour (Murray-Watson et al., 2023) driven by information on factors affecting stakeholder opinions (Garcia-Figuera et al., 2021; Exilien et al., 2024). Current HLB contingency plans in Portugal and Spain (DGAV, 2021; BOE, 2023) include a buffer zone surrounding the infested area within which intensive surveys and coordinated insecticide sprays should be applied. Modelling could again be used, to optimise the size of the buffer zone and the type of surveillance to be applied within it, to provide quantitative support for contingency plans.

Slowing transmission by heavily controlling vector populations with additional insecticide is effective in our model (Figs. 7(D) and 8(D)), as it has been in Brazilian citriculture (Bassanezi et al., 2020). However, our model’s representation of vectors might overstate efficacy. We do not model vector population dynamics, and so in turn we assume psyllid densities are immediately/simultaneously reduced in managed regions, ignoring difficulties of attaining such area-wide control (Galvañ et al., 2023). We also assume psyllid populations can be reduced by up to 99%, without considering the frequency of sprays required, nor the expense, nor risks of insecticide resistance (which is now emerging in Brazil). Requisite chemical doses to achieve such reductions seem unlikely to be consistent with EU regulations (Lázaro et al., 2021). While Regulation (EU) 1107/2009 does permit emergency authorisation when a pest cannot be controlled by other means (European Union, 2019), this applies for only a limited time. Indeed, active ingredients currently labelled for citrus pests in the EU are less effective than those referred to for model parameterisation; most chemicals in Qureshi et al. (2014) are no longer marketed. In practice, it is also particularly difficult to protect flush (i.e., young) leaf tissue favoured by psyllids and implicated in transmission (Cifuentes-Arenas et al., 2018), since the most commercially attractive (i.e., cheapest) insecticides are not systemic and do not cover rapidly growing tissue. However, controlling flushing frequency via selective pruning might provide partial mitigation (Matias et al., 2023).

Despite many unavoidable uncertainties, modelling provides the only mechanism to understand how HLB might spread in the EU, and to answer questions surrounding the best approach to detect and control any outbreak. We conclude that the most efficient management strategy would include early detection and intensive roguing to remove inoculum, alongside other measures to slow spread, particularly enhanced pest management to control psyllids. However, even very effective management will not eradicate any epidemic, and ensuring engagement from growers is essential. Following first detection, the focus will shift to sustaining the citrus industry for the longest possible time in the face of HLB (Bassanezi et al., 2020). This is another area in which modelling can play a prominent role.

## Acknowledgements

The work was supported by Pre-HLB (Preventing HLB epidemics for ensuring citrus survival in Europe), Grant 817526 from the European Union Horizon 2020 program. Additionally, T.M. acknowledges support from FCT for 2020.07798.BD (doi: 10.54499/2020.07798.BD), and T.M and A.D. jointly acknowledge support from MED for UIDB/05183/2020 (doi: 10.54499/UIDB/05183/2020) and from UIDP/05183/2020 (doi: 10.54499/UIDP/05183/2020) and from CHANGE for LA/P/0121/2020 (doi: 10.54499/LA/P/0121/2020). J.B.-M. and J.A.P. also jointly additionally acknowledge support from FCT/MCTES (PIDDAC) for CIMO, UIDB/00690/2020 (doi: 10.54499/UIDB/00690/2020) and UIDP/00690/2020 (doi: 10.54499/UIDP/00690/2020); and SusTEC, LA/P/0007/2020 (doi: 10.54499/LA/P/0007/2020).

## Author contributions

J.E. and N.J.C. designed the modelling framework and parameter estimation approach, and selected scenarios to test using the fitted model with input from E.L., A.V. and S.P. in identifying scenarios to test. J.E. developed and tested the computational code. E.L., B.D., T.M., A.D., J.B.-M. and J.A.P. provided or processed citrus host and/or psyllid data. J.E. and N.J.C. wrote the manuscript, with input from all co-authors.

## Data availability

Code and data are on GitHub https://github.com/DrJREllis/HLBinEurope.

## Supporting Information

- S1 Supporting Methods
- S2 Supporting Results
- S3 Supporting Videos

## Open Access

For the purpose of open access, the author has applied a Creative Commons Attribution (CC BY) licence to any Author Accepted Manuscript version arising from this submission.

