## Supplementary Information for "Developing epidemiological preparedness for a plant disease invasion: modelling citrus huánglóngbìng in the European Union"

Supporting Information consists of the following sections.

- S1 Supporting Methods
- S2 Supporting Results
- S3 Supporting Videos
- References

### S1 Supporting Methods

#### S1.1 Mapping citrus and climatic suitability

##### S1.1.1 Citrus densities

We used publicly-available official agricultural statistics and census data for Spain and Portugal (Ministerio de Agricultura, Pesca y Alimentación, 2000-2020; Instituto de Financiamento de Agricultura e Pescas, 2021; Eurostat, 2016) to map densities of residential and commercial citrus across the Iberian peninsula, following the procedure described in full in Galvañ et al. (2023). Data crossing and layer/raster design was done with QGIS 3.12.3 (QGIS.org, 2020). For each 1km × 1km cell, $i$, this led to estimates of the densities of commercial ($h_i^c$) and residential ($h_i^r$) citrus, i.e., the maps shown in Fig. 1(B) of the main text.

In converting host densities to numbers of citrus host units for use in our epidemiological model, $H_i^c$ and $H_i^r$, we used different discretisations for commercial and residential citrus. The underpinning idea was to allow our model to capture within-cell changes of infection densities in both classes of citrus over appropriate ranges, since typical values of $h_i^c$ and $h_i^r$ tended to be rather different (see also Fig. 1(C) in the main text). A single commercial host unit was taken to correspond to an area of 100m × 100m (i.e., 1 ha), meaning those cells entirely covered by commercial citrus would contain 100 commercial host units. A single residential citrus host unit was instead taken to correspond to an area of approximately 32m × 32m (0.1 ha), meaning there was an upper limit of $1,000$ residential host units per cell. The largest number of residential host units in any cell across the whole of Spain and Portugal was in fact 71 (out of a potential maximum of $1,000$), corresponding to a 1km × 1km (i.e., 100 ha) cell with 7.1 ha of trees in residential/municipal settings.

Our mapping procedure allowed the density of commercial citrus in cell $i$ (i.e., $h_i^c$) to be partitioned into three classes: typical (actively cultivated, with insecticides sprayed), abandoned (not cultivated) and organic (only cultural control). When modelling pest management as applied by commercial growers, we assumed no insecticides were sprayed in abandoned and organic citrus, by taking the effect of management in cell $i$ to be

$$m_i = 1 - m\left(1 - \frac{h_i^a + h_i^o}{h_i^c}\right), \quad (1)$$

where $m$ is a constant that determines the effectiveness of pest management ($0 \leq m \leq 1$; we selected $m = 0.9$ by default with reference to Qureshi et al. (2014)), and where $h_i^a$ and $h_i^o$ are the proportions of cell $i$ with abandoned and organic commercial citrus, respectively.

##### S1.1.2 Climatic suitability

The response of psyllid population density to climate was modelled using daily weather data for Spain and Portugal, obtained from the ERA5-Land dataset (Copernicus Climate Data Store, 2019). Following previous studies of AfCP (Catling, 1973), a day was considered "suitable" whenever the vapor pressure deficit was less than 34.5 mbar (Paiva et al., 2020), and the minimum temperature was greater than 10°C. The climate suitability, $w_i$ (mapped in Fig. 1(A) in the main text) is simply the proportion of days with suitable conditions taken over the entire 2009 to 2018 period.

#### S1.2 Dispersal kernels

As described in the main text, we separately capture two scales of dispersal: local (i.e., short-distance) and long-distance. In turn this requires two dispersal kernels.

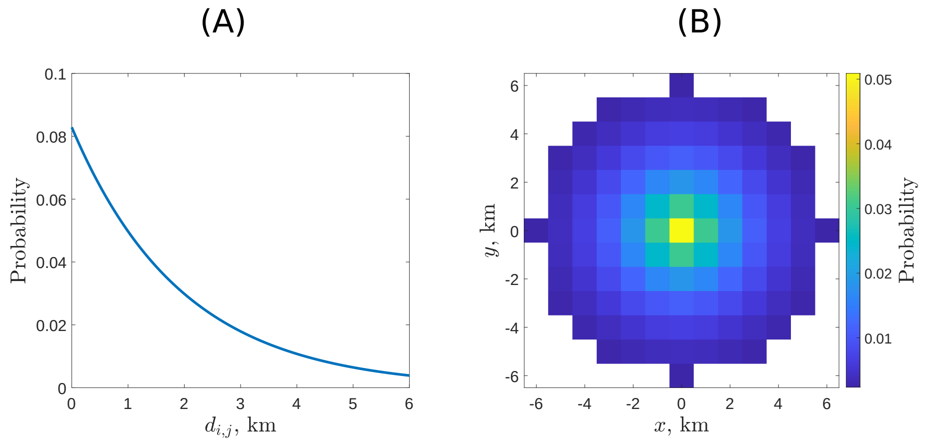


**Figure S1: Local dispersal kernel.** (A) Shown as a 1D function of $d_{i,j}$, the distance between the centres of cells $i$ and $j$. (B) Shown as a normalised probability density function in 2D (Eqn. 2). Note that the kernel is truncated so that (local) dispersal is not permitted between cells with centres more than $d_{max} = 6$km apart. Psyllid dispersal over longer distances is accounted for via the long-distance dispersal kernel (Fig. S2).

###### S1.2.1 Local dispersal

The local dispersal kernel (Fig. S1) between a pair of cells $i$ and $j$ is

$$K_{ij}^{loc} = \begin{cases} A\exp\left(-d_{ij}/\alpha_{loc}\right) & \text{for } d_{ij} \leq d_{max} \\ 0 & \text{otherwise} \end{cases}, \tag{2}$$

where $d_{ij}$ is the distance between centres of cells $i$ and $j$, $\alpha_{loc} = 1.96$km is the scale parameter for short-ranged dispersal (chosen by reference to Nguyen et al. (2023)), $d_{max} = 6$km is the maximum range over which local dispersal occurs, and $A$ is a normalising constant selected to ensure $K_{ij}^{loc}$ is a valid probability mass function, i.e.,

$$A = \left(\sum_{(k,l)\in\Omega} \exp\left(-\sqrt{k^2+l^2}/\alpha_{loc}\right)\right)^{-1}, \tag{3}$$

where $\Omega$ is the subset of $\mathbb{Z}^2$ corresponding to cells within distance $d_{max}$ of the origin, i.e.,

$$\Omega = \left\{(k,l)|k \in \mathbb{Z}, l \in \mathbb{Z}, \sqrt{k^2+l^2} \le d_{max}\right\}. \tag{4}$$

We limited local dispersal to cells within a distance of $d_{max} = 6$km for computational efficiency. However, we did exploratory runs to verify this had no marked effect on our results (for the value of $\alpha_{loc} = 1.96$km as used in our fitted model, only a very small proportion of local dispersal would be further than 6km; Fig. S1).

##### S1.2.2 Long-distance dispersal

Our long distance kernel (Fig. S2) is a $t$-distribution with 3 degrees of freedom, i.e.,

$$K^{ld}(d) = \frac{\Gamma(2)}{\Gamma(3/2)\sqrt{3\pi}\alpha_{ld}}\left(1+\frac{1}{3}\frac{d^2}{\alpha_{ld}^2}\right)^{-2}, \tag{5}$$

where $d$ is the distance over which dispersal occurs, $\alpha_{ld} = 130$km (chosen by reference to Benhadi-Marín et al. (2020)) is the scale parameter for long distance dispersal, and $\Gamma(\cdot)$ is the Gamma function (i.e., the function defined as $\Gamma(z) = \int_0^\infty t^{z-1}e^{-t}dt$).

As described in the main text, we use the "particle-emission" formulation of dispersal originally introduced in the model of Meentemeyer et al. (2011). This is because – at the spatial scales considered here – simulating long-distance psyllid dispersal by accounting for ever-changing forces of infestation on each uninfested cell across the entire landscape in our Gillespie algorithm simulations would be computationally intractable. We therefore invert the problem, and model rates of long-range dispersals emanating from colonised cells. This requires us to sample dispersal distances from the probability distribution in Eqn. 5 whenever a long-distance dispersal occurs; we did this by inversion sampling.

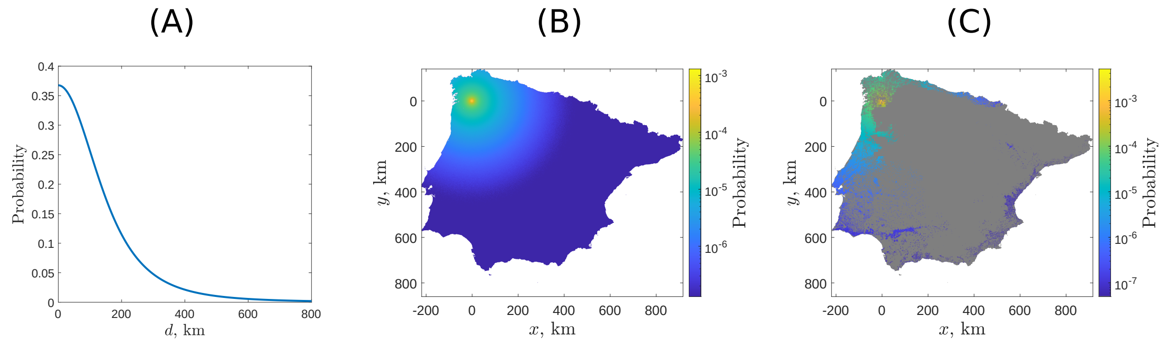


**Figure S2: Long-distance dispersal kernel.** (A) Normalised (1D) probability density function used for long distance dispersal (Eqn. 5). (B) Relative probabilities of dispersal to different locations when originating from a point in Galicia, North-West Spain. (C) Probabilities of arriving specifically in cells that contain citrus host units, starting from the same point as in (B). The colour gradient in (B) and (C) is on a log scale.

#### S1.3 Model fitting

##### S1.3.1 Psyllid within-cell colonisation and dispersal

**Sources of data.** We used AfCP presence data from surveys in Portugal and Spain up to 2021 to calibrate psyllid dispersal in our model (Benhadi-Marín et al., 2022). AfCP was first located in Galicia in north-western Spain in 2014 (Cocuzza et al., 2017; Perez-Otero et al., 2015), and subsequently spread along the western coast of Portugal and the northern coast of Spain. However, starting in autumn 2019, the parasitoid *Tamarixia dryi* (Waterson) began to be released in multiple locations in the infested area, greatly reducing psyllid populations (Molina et al., 2021; Duarte et al., 2024). For the purposes of parameterisation, we therefore assume that the vast majority of psyllid dispersal had already occurred by 2019.

**Within-cell psyllid colonisation.** It is clear that psyllid population growth is linked to seasonal cycles of citrus flush growth (Udell et al., 2017), and it may take several of these cycles for the psyllid population to reach a stage where it has largely colonised 1km × 1km grid cell. We also note that in our model all citrus of a particular type within each cell is either psyllid infested or not, and some lag to reach full population density must be required. However, despite interrogating a number of sources collating small-scale AfCP population dynamics data (Catling, 1973; Samways and Manicom, 1983; van den Berg et al., 1991; Tamesse and Messi, 2004; Cook et al., 2014), we were not able to find suitable sources of information to fix an appropriate time scale. We therefore set the average time of colonisation to be one year ($\eta$ = 1/365 $d^{-1}$), noting that this was somewhat arbitrary. However, as described below, the method we adopted to find values of $\delta$ and $\zeta$ accounts for the arbitrariness of this choice.

**Finding dispersal parameters.** The parameters $\delta$ (dispersal rate of psyllids) and $\zeta$ (proportion of long distance dispersal) were fitted to AfCP presence data in the Iberian Peninsula (Benhadi-Marín et al., 2022). The aim of model fitting was to select the pair of values $\delta$-$\zeta$ which caused the results of our model to most closely match the survey data, a single “snapshot” of the extent of the infestation in 2019 as reported in 2021 (Fig. S3).

For different pairs values of $\delta$ and $\zeta$, we used our model to simulate psyllid population dynamics and spread following the introduction of the psyllid into ten 1km × 1km cells in the near vicinity of where it was first reported in Galicia. We therefore assumed there was already a small population of vectors in 2013, one year before AfCP was first detected in Galicia. Our simulations ran over a 6 year period, corresponding to the time over which the psyllid was able to spread freely and before the parasitoid *Tamarixia dryi* was released in Portugal. Simulations were repeated 300 times for each pair of parameters.

A "Log Score", representing the log likelihood of the survey data according to our model, was then calculated as a measure of model prediction accuracy. This was done by assuming the presence/absence status of each cell in the survey data corresponded to the result of a single trial in a series of independent Bernoulli trials, where the probability of vector presence in a cell was estimated empirically as the

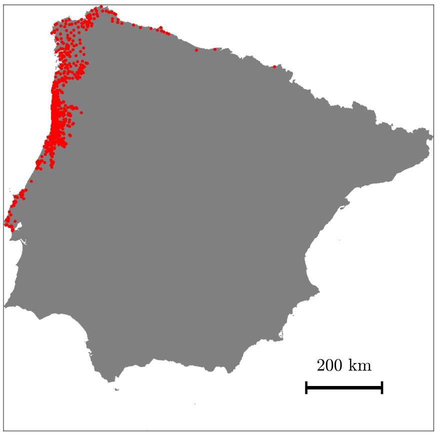


**Figure S3: AfCP presence in the Iberian peninsula as of 2021**. Red dots indicate locations where AfCP had been found by 2021 (Benhadi-Marín et al., 2022), following first detection in Galicia (northwestern Spain) in 2014 (Perez-Otero et al., 2015).

fraction of the 300 simulations for which the cell was invaded for the pair of parameters in question. In particular, the Log Score for parameter values $\delta$ and $\zeta$ is

$$L(\delta,\zeta) = \sum_j \big(Z_j \ln(p_j) + (1 - Z_j)\ln(1 - p_j)\big), \tag{6}$$

where $Z_j$ is the binary variable indicating vector presence in cell $j$ in the survey data, and $p_j$ is the probability of the vector dispersing to that grid cell as estimated over 300 simulations for the pair of parameters in question. Large values of $L(\delta,\zeta)$ indicate the model using the pair of parameters $(\delta,\zeta)$ is a good fit to the survey data, meaning that maximising $L(\delta,\zeta)$ was the aim of model fitting.

**Upscaling the survey data.** Before doing the calculation in Eqn. 6, the simulated and surveyed vector presence data at the 1km × 1km scale were converted to a coarser 20km × 20km grid. This was done so that small differences in the exact location of the psyllids

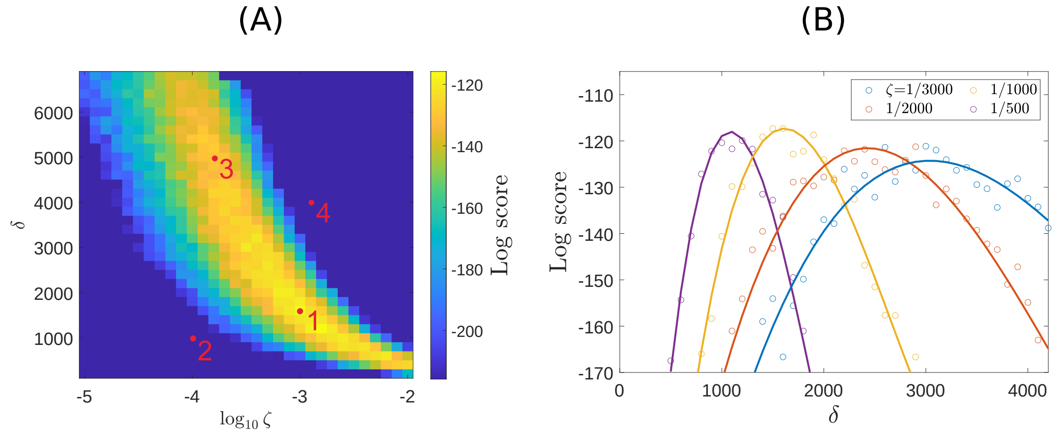


**Figure S4: Parameter scan for dispersal parameters.** (A) Log Scores (Eqn. 6) presented in 2D with varying dispersal rate ($\delta$) and proportion of dispersals that are long distance ($\zeta$) and (B) multiple 1D plots with fixed $\zeta$ and varying $\delta$. For each fixed $\zeta$ there is a maximum, but there is no clear maximum in the 2D space. The dark blue spaces on the left figure show regions where the Log Score was less than the maximum by 100 or more. See also Figs. S5 and S6, corresponding to the points marked 1 and 2 to 4 respectively in (A), for examples of the predicted vector dispersal that the Log Score is evaluating.

in individual runs of our model would not be penalised by the model fitting. Without such an upscaling procedure, if an individual run of our model predicted the vector would spread to a specific 1km × 1km cell on the west coast of Portugal, but, though the vector has been found in many locations nearby, it was not found in the survey data in precisely that location (since, for example, that location was not inspected), this would adversely affect our assessment of the model's parameterisation. We tested different grid sizes for our upscaled grid, ranging from 5km × 5km to 50km × 50km, and found that while using a coarser grid was important to attain reliable results, the particular size of the coarser grid used did not substantially affect our inferences concerning vector dispersal.

**Results of model fitting.** The Log Scores for the vector dispersal parameterisation were calculated using Eqn. (6), for a range of values of $\delta$ and $\zeta$, the results of which are visualised in Fig. S4. Unsurpris-

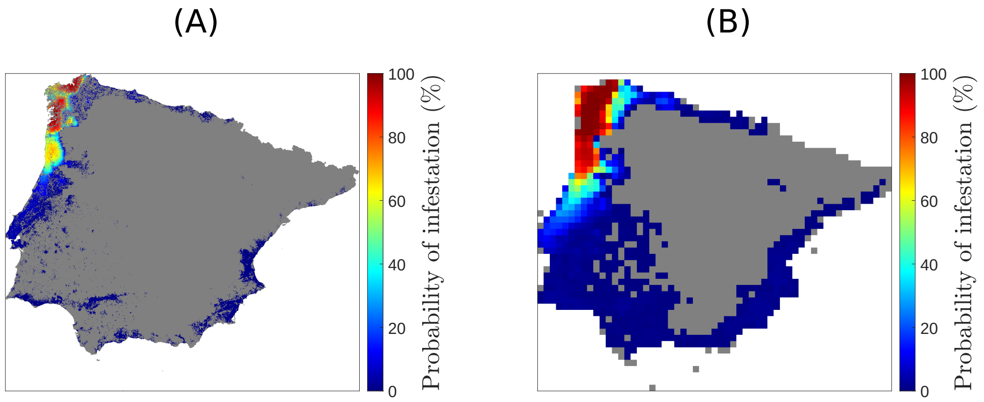


**Figure S5: Spread of AfCP using the best fitting model parameters.** Maps showing the simulated probability after 6 years of spread of AfCP in Iberia, following introduction in northwestern Spain, with optimal parameter $\delta = 1600\text{d}^{-1}$ and $\zeta = 10^{-3}$, resulting in Log Score $= L(\delta, \zeta) = -117$ (Point 1 on Fig. S4). (A) the probability of the vector being present in each original 1km × 1km grid cell and (B) the probability of the vector being anywhere within each coarse 20km × 20km grid cell. The Log Score compares predicted spread to the locations from the data set where AfCP has previously been detected (Fig. S3). Although more of the map is coloured than where the vector is present in the data set, note dark blue indicates a low probability (< 10%) that the vector will travel to a particular locations.

ingly, since essentially a single data point (the extent of the infestation in 2019 as reported in 2021) is being used to identify two parameters, there is no single clear maximum of the Log Score in the parameter space. However there is a region in which $1,000 < \delta < 2,000$ ($\delta$ has units $\text{d}^{-1}$), and $-3.5 < \log_{10}\zeta < -2.5$, which has slightly higher Log Scores than pairs of parameters with larger $\delta$ and smaller $\zeta$ (the yellow region in Fig. S4).

Examples of the coarser 20 km × 20 km maps used to calculate the Log Scores are in Figs. S5 and S6. In the maps with higher scores, the model predicts a high probability that the psyllids will spread down the western coast of Portugal, with a probability that gradually decreases at locations more distant from northwestern Spain and Portugal, with low but non-zero probabilities that the vector will spread widely around Iberia. In cases with small $\delta$ and $\zeta$ (to the left or below the yellow region in Fig. S4), the vector has a low probability of

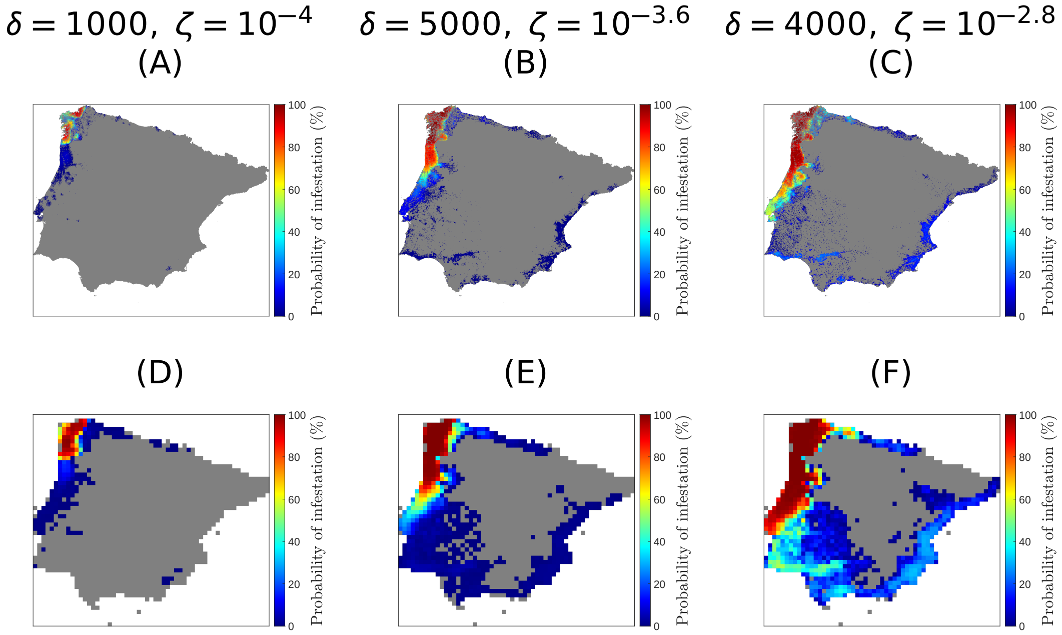


**Figure S6: Spread of AfCP using sub-optimal model parameters.** Maps showing the probability of the spread of AfCP in Iberia using sub-optimal dispersal parameters on the original 1km × 1km grid (A, B, C) and the coarse 20km × 20km grid (D, E, F). (A, D) show a parameter choice which spreads too slowly (Log Score $= L(\delta,\zeta) = -263$; Point 2 on Fig. S4), (B, E) show a choice with a larger $\delta$ and smaller $\zeta$, which performs similarly well to the best fitting parameters (*cf.* Fig. S5) (Log Score $= L(\delta,\zeta) = -130$; Point 3 on Fig. S4), (C, F) show a parameter choice that results in spread of the vector that is too fast (Log Score $= L(\delta,\zeta) = -460$; Point 4 on Fig. S4).

spreading very far outside the region to which it was introduced initially (Fig. S6(A) and Fig. S6(D)). In cases with large $\delta$ and $\zeta$ (to the right or above the yellow region in Fig. S4), the vector is likely to spread widely, often reaching the coast of eastern Spain (Fig. S6(C) and Fig. S6(F)).

To determine which parameter pair to take in our fitted model, we selected a small number of regularly-spaced fixed values of $\zeta$ and for each plotted the Log Score against $\delta$ (Fig. S4). There was a value of $\delta$ at which the Log Score was maximum for each chosen $\zeta$. From these we selected the value of $\zeta$ that peaked at the highest score, which gives $\zeta = 10^{-3} = 1/1000$ and the corresponding maximum at

$\delta = 1600\mathrm{d}^{-1}$. The probability of infestation for this pair of parameters is mapped in Fig. S5.

As described above, to account for the colonisation rate of the vector, we used a value of $\eta = 1/365\mathrm{d}^{-1}$ (i.e., 1 per year). A different value of $\eta$ results in different values of $\delta$ and $\zeta$, but the overall quality of the fitted model is not affected, as the fitting ensures that the vector spreads at the same overall rate. This is because the comparison between simulated and real data only uses populations that have fully colonised, so for example, if the colonisation rate were slower, the vector would have to arrive in a location earlier for it to reliably have colonised that location by 2019. This would therefore necessitate a faster rate of dispersal, which would be achieved by the fitting.

##### S1.3.2 Infection rate parameters

**Overview.** To fit HLB infection rates, we calibrated our model against Mastin et al. (2020), a model of the spread of (*C*Las) HLB in Florida. Even if the differences in model structure could be systematically corrected between our model and the Mastin et al. (2020) model, parameter estimates cannot be used directly due to the differences in the host landscape. In particular, in Florida, vast areas of cultivation cover many tens of square kilometres, which is not the case in Spain or Portugal (see Fig. 1(A) in main text). We therefore instead compare the spread of infection in both models when applied to the same host landscapes, which were randomly generated to approximate a host distribution similar to those in areas of high commercial citrus in Spain. The parameters to be fitted were the overall infection rate, $\beta$, and the proportion of infections that occur within cell, $\rho$. This was again done by scanning over pairs of parameter values, attempting to find pairs for which the results of our model most closely match a measure of “ground truth”.

**Sources of data.** As described above, here "ground truth" corresponds to the results of the Mastin et al. (2020) model on landscapes more typical of citrus cultivation in Iberia. In particular, we sampled regions of high density commercial citrus across the Iberian peninsula to produce a set of 100 exemplar 15km × 15km maps upon which to simulate disease spread. To do this, we sampled 3km × 3km blocks of commercial citrus from the entire map where the minimum density was $h_i^c \geq 0.02$ and randomly patched them together in groups of 5 to create new landscapes.

**Finding infection rate parameters.** The Mastin et al. (2020) model was run 100 times on each map for 10 years and the mean proportion of cells and proportion of citrus infected across each 15km × 15km landscape calculated at the end of each year. These metrics were also extracted from ensembles of simulation runs of our model, each time using a particular pair of values $\beta$ and $\rho$, and averaging over 100 model runs per host landscape. The measure of deviation between the two models was taken to be the squared difference in the simulated proportions of infectious cells and citrus between the two models, summed over all 10 years of spread over all maps, i.e.,

$$E(\beta,\rho) = \sum_{y=1}^{10}\left(\overline{I+C} - \overline{M}\right)^2 + \sum_{y=1}^{10}\left(\overline{1_{I+C}} - \overline{1_M}\right)^2 \qquad (7)$$

where $\overline{I+C}$ is the mean proportion of infectious citrus in our model, $\overline{M}$ is the mean total amount of infected citrus in the Mastin et al. (2020) model, and $\overline{1_{I+C}}$ and $\overline{1_M}$ are the mean proportions of cells that have at least one unit of infected citrus for our model and the Mastin et al. (2020) model, respectively.

**Results of model fitting.** The mean squared difference between the proportion of infectious citrus units in each model and the proportion of cells with infectious citrus as calculated using Eqn. 7 as $\beta$ and

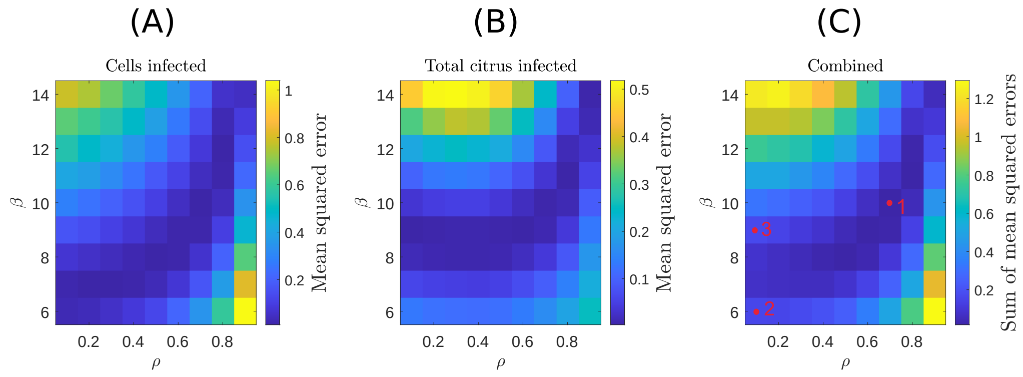


**Figure S7: Finding infection rate parameters.** Mean squared error between the results of our model and Mastin et al. (2020) model for each pair of $\beta$ and $\rho$ between (A) the proportion of cells infected, (B) the proportion of citrus infected and (C) the sum of both. The overall infection rate is given by $\beta$ and the proportion of infections that remain within the cell that they originate is given by $\rho$. See Figs. S8 and S9 for examples comparing disease progress curves corresponding to Points 1-3.

$\rho$ are varied are shown in Fig. S7. Comparisons between average disease progress curves when using the different parameters are shown in Fig. S8 and S9.

In cases with small values of $\rho$, in which the majority of infections originating from infected citrus are dispersed to nearby cells instead of within the same cell, the optimal infection rate, $\beta$, for the two measures of infection do not correlate. As demonstrated in Fig. S9, when $\beta = 6\text{d}^{-1}$, the proportion of cells infected is similar to that in the Mastin et al. (2020) model, however because the overall infection rate is low, the proportion of citrus infected remains low. Conversely, when $\beta = 10\text{d}^{-1}$, the proportion of citrus infected is a good fit to the Mastin et al. (2020) model, but when $\rho = 0.1$, there is too much infection occurring outside of the cell, so that the proportion of cells infected is too high.

It can be seen in Fig. S7 that the optimum of the two measures coincide at $\beta = 10\text{d}^{-1}$ and $\rho = 0.7$. This pair of values – which gives the overall minimum squared difference and produces simulations that result in a proportion of cells with infected citrus and an overall proportion of infected citrus that is similar to the Mastin et al. (2020)

model (see Fig. S8) – is therefore the result of our model fit.

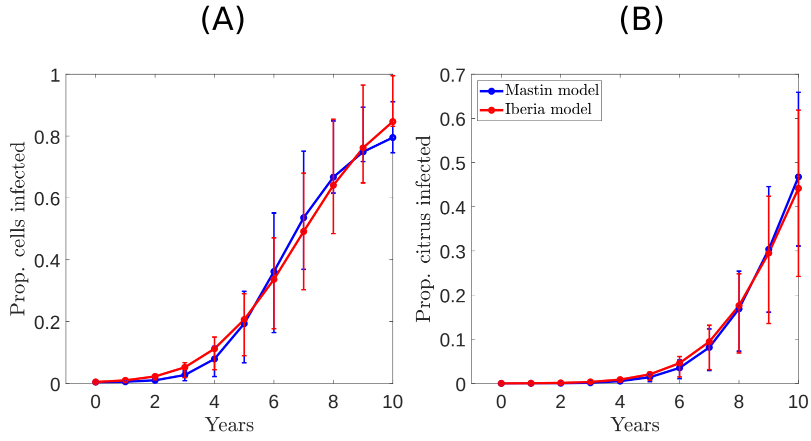


**Figure S8: Model comparison for best fitting infection rate parameters.** Comparison of progression curves for the mean and interquartile range of (A) the proportion of cells and (B) total citrus infected when using the parameters that give the minimum combined mean squared difference (see Point 1 in Fig. S7): $\beta = 10\text{d}^{-1}$ and $\rho = 0.7$.

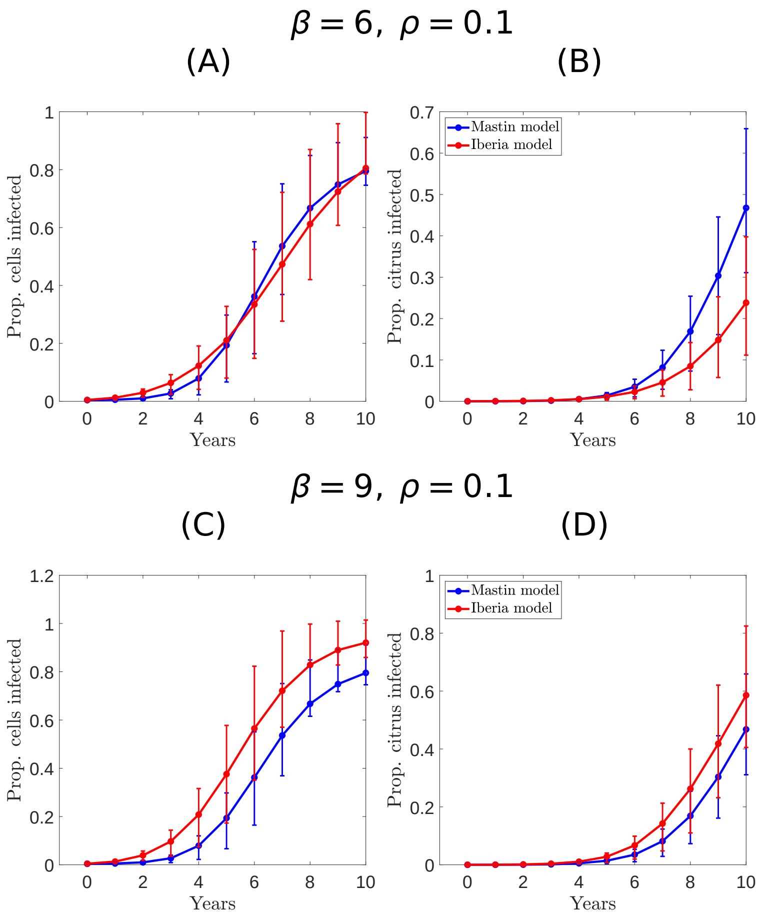


**Figure S9: Examples of model comparisons for poorly fitting infection rate parameters.** Comparison of progression curves for the mean proportion of cells and mean total citrus infected and interquartile range from the two models with suboptimal values of $\beta$ and $\rho$. In (A, B), corresponding to Point 2 in Fig. S7, there is a relatively good fit for the proportion of cells infected, but the proportion of citrus units infected is under-predicted. In (C, D), corresponding to Point 3 in Fig. S7, the proportion of citrus is a good fit, but the proportion of cells infected is over-predicted.

### S2 Supporting Results

#### S2.1 Additional results for Region A

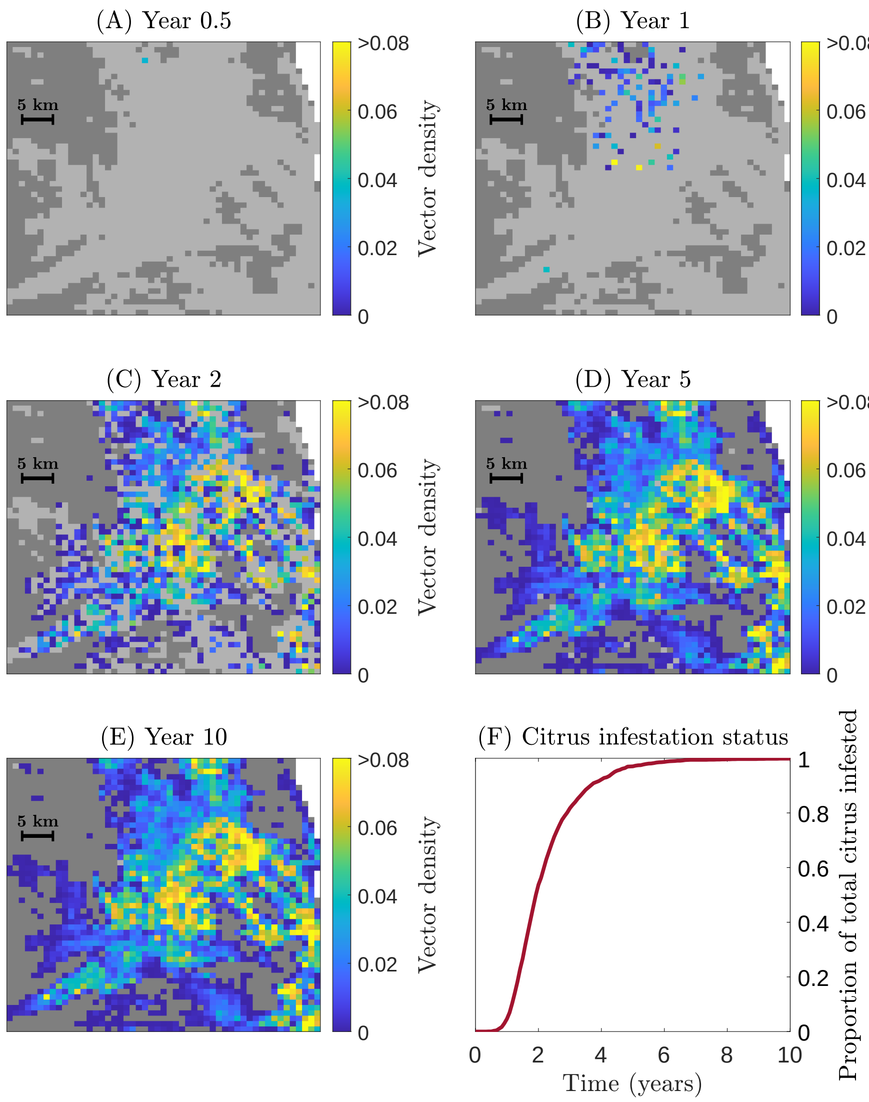


**Figure S10: Spread of the vector in a single simulation in Region A (Valencia).** Underpinning spread of the vector in the single simulation visualised in Fig. 3 in the main text. (A) Maps show the measure of vector density within each cell ($V^r + V^c$). Only 10 years of spread are shown (*cf.* 20 years for the maps showing pathogen spread in Fig. 3 in the main text) because the vector has already fully colonised the region within 10 years. (F) shows the proportion of citrus that is vector infested as a function of time. See also S3 Supporting Videos (Video 1), which also shows initial exposure in light grey.

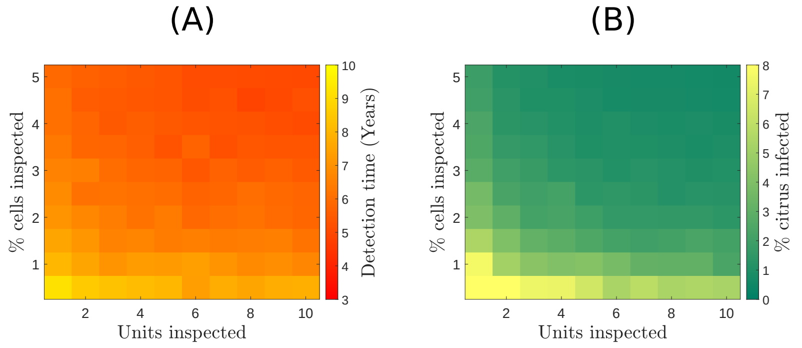


**Figure S11: Effectiveness of surveillance strategies in Region A (Valencia).** Responses to varying the number of host units inspected per surveyed cell ($x$-axis) and the percentage of cells inspected ($y$-axis) (see also Fig. 5 in the main text). (A) Mean time of detection and (B) Mean percentage of total citrus infected at the time of detection. The interval between inspections is $\Delta = 1$ year and the probability of detection is $p = 0.5$.

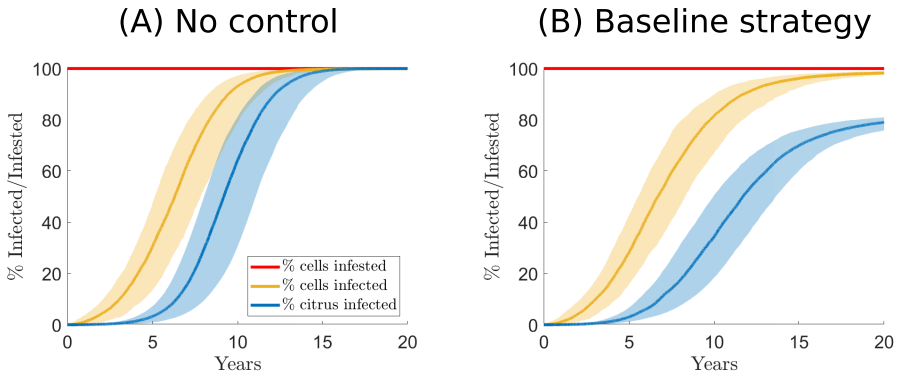


**Figure S12: Effect of the baseline control strategy on the spread of the pathogen in Region A (Valencia).** The vector is widespread at the time of infection, hence the percentage of cells infested by the vector is at 100% (red), other curves show the percentage of cells with at least one infected unit of citrus (yellow) and the percentage of all citrus infected (blue), without any control strategy (Panel (A)) *versus* with the baseline control strategy (Panel (B)). In panel (B), infected citrus includes those that have been infected and subsequently removed. In all cases, the pathogen is introduced into the same cell at $t = 0$; the cell initially infected is chosen at random. Shaded regions show 95% prediction intervals from 200 simulation results.

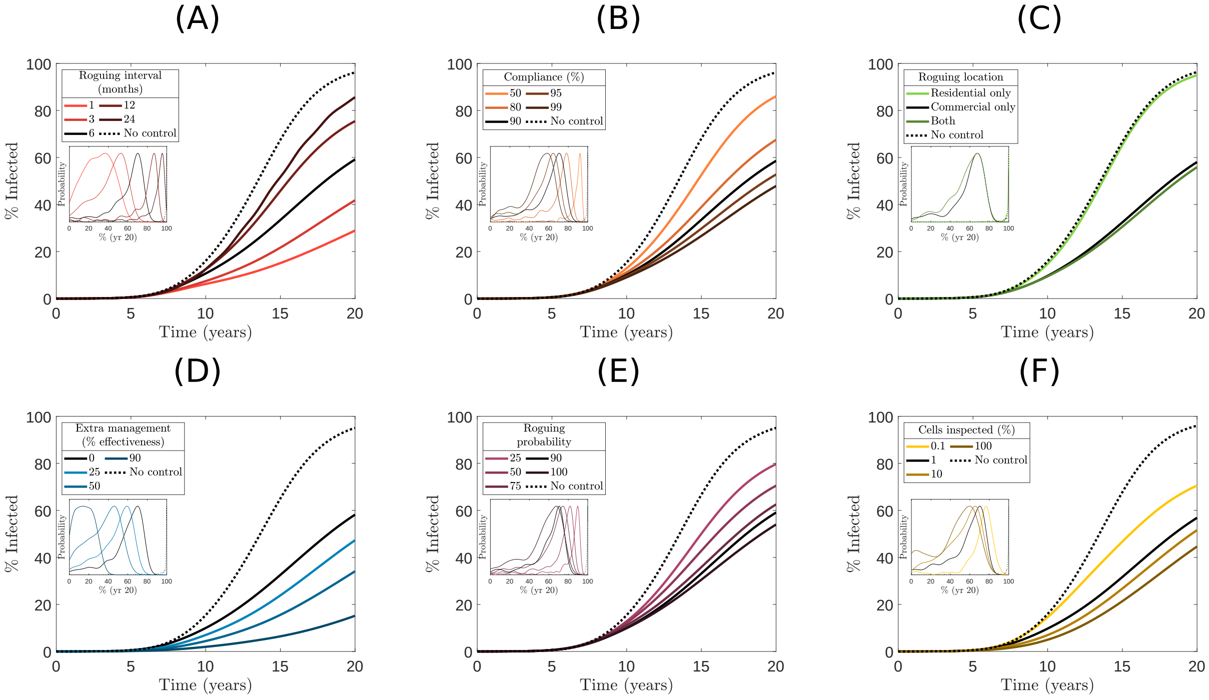


**Figure S13: Management scenarios in Region A (Valencia) when the vector is spreading.** Mean proportions of infected or removed citrus ($E + C + I + R$) over time when varying a single parameter from the baseline case (Fig. 6 and Table 2 in the main text), but when the vector is introduced to the same cell as the pathogen at $t = 0$. This figure is therefore analogous to Fig 7. in the main text, but shows the effect of the vector also invading on our results. Aspects varied in each panel: (A) roguing interval, (B) compliance, (C) types of citrus rogued, (D) increases to the pest management parameter, $m*$, (E) roguing probability, and (F) proportion of cells inspected for early detection. Inset graphs show probability distributions of the proportion of infected or removed citrus after 20 years (normalised to have the same maximum for ease of visualisation). Black lines show results using the baseline parameterisation; dotted lines show results with no control. Averages and probability distributions calculated from 200 simulations per parameter combination.

#### S2.2 Additional results for Region B

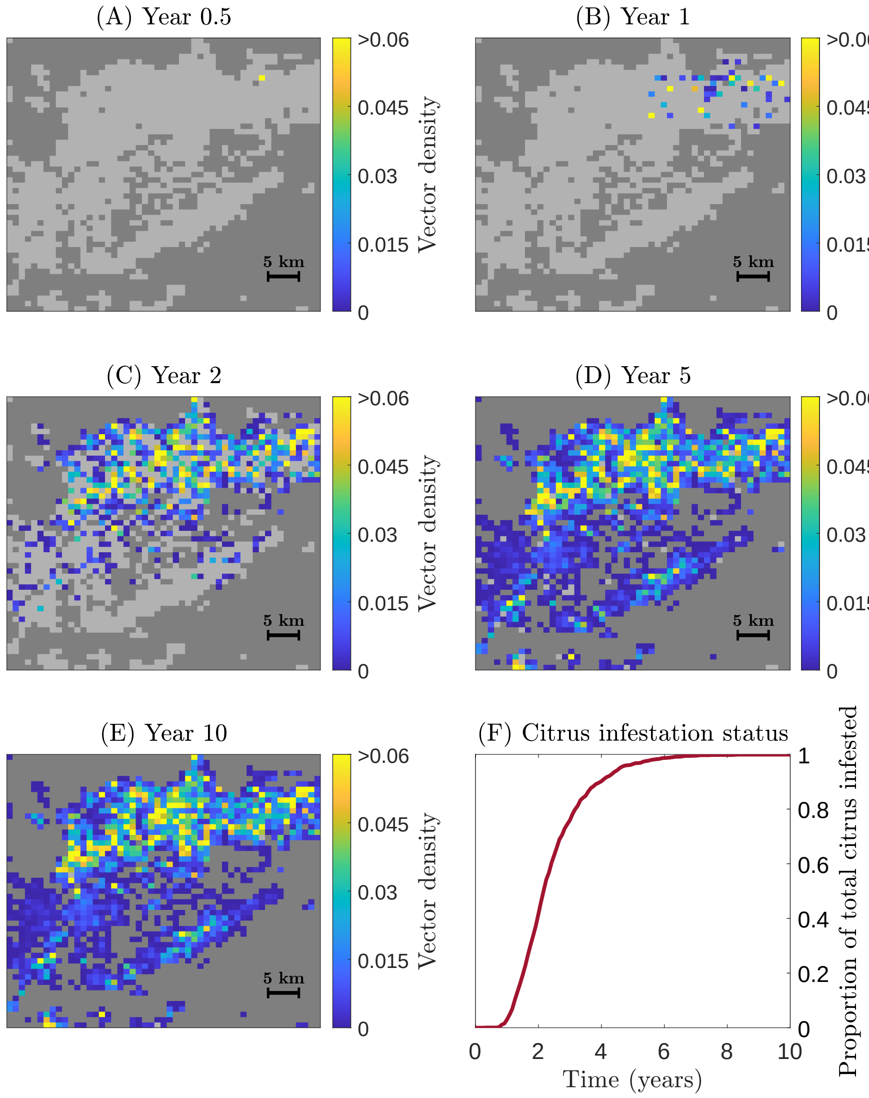


**Figure S14: Spread of the vector in a single simulation in Region B (Andalusia).** The vector was introduced to commercial citrus in a single cell at $t = 0$; marked with the yellow square in panel (A). Maps show the measure of vector density within each cell ($V^r + V^c$). Again the entire region is fully colonised with 10 years. Panel (F) shows the proportion of citrus that is infested over the entire region. See also S3 Supporting Videos (Video 4), which also shows initial exposure in light grey.

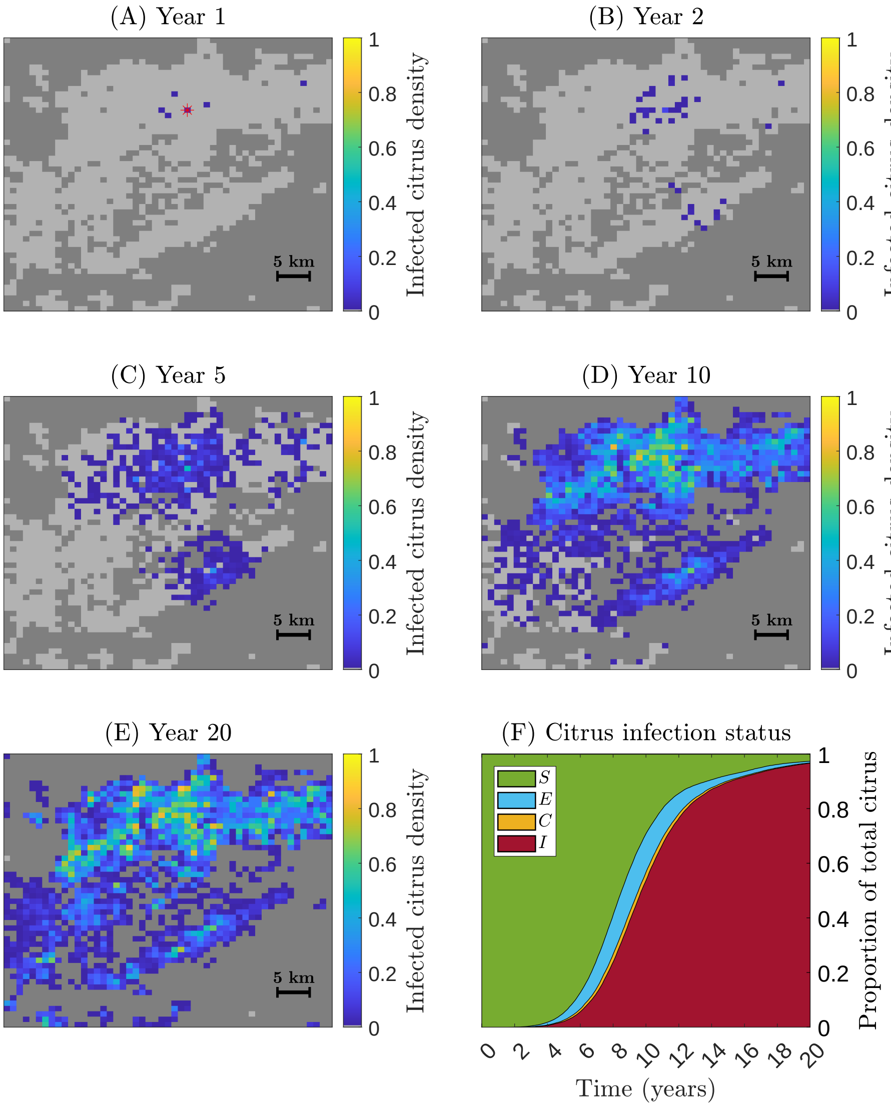


**Figure S15: Spread of the pathogen in a single simulation in Region B (Andalusia).** A single citrus host unit exposed to the pathogen was introduced to a single cell at $t = 0$. Maps show the density of infected citrus host units ($E + C + I$) within each cell. The disease progress curve (F) shows the proportion of all citrus over the entire region in each epidemiological compartment, (S)usceptible, (E)xposed, (C)ryptic and (I)nfected. See also S3 Supporting Videos (Video 5).

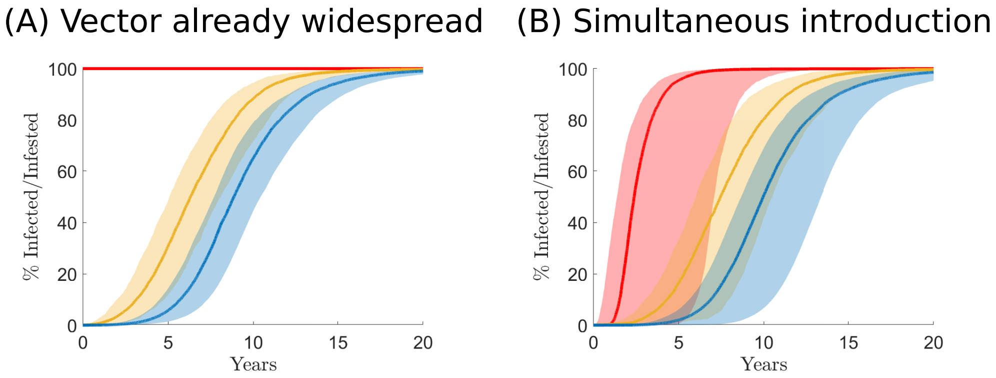


**Figure S16: Effect of prior invasion by the vector on the spread of the pathogen in Region B (Andalusia).** The percentage of cells infested by the vector (red), with at least one infected unit of citrus (yellow) and the percentage of all citrus infected (blue), when the vector is already fully established throughout the region of interest (Panel (A)) *versus* when the vector and pathogen are introduced simultaneously (Panel (B)). In all cases the pathogen is introduced into the same cell at $t = 0$; for the simulations shown in Panel (B), the cell initially infested by the vector is chosen at random. Shaded regions show 95% prediction intervals from 200 simulation results.

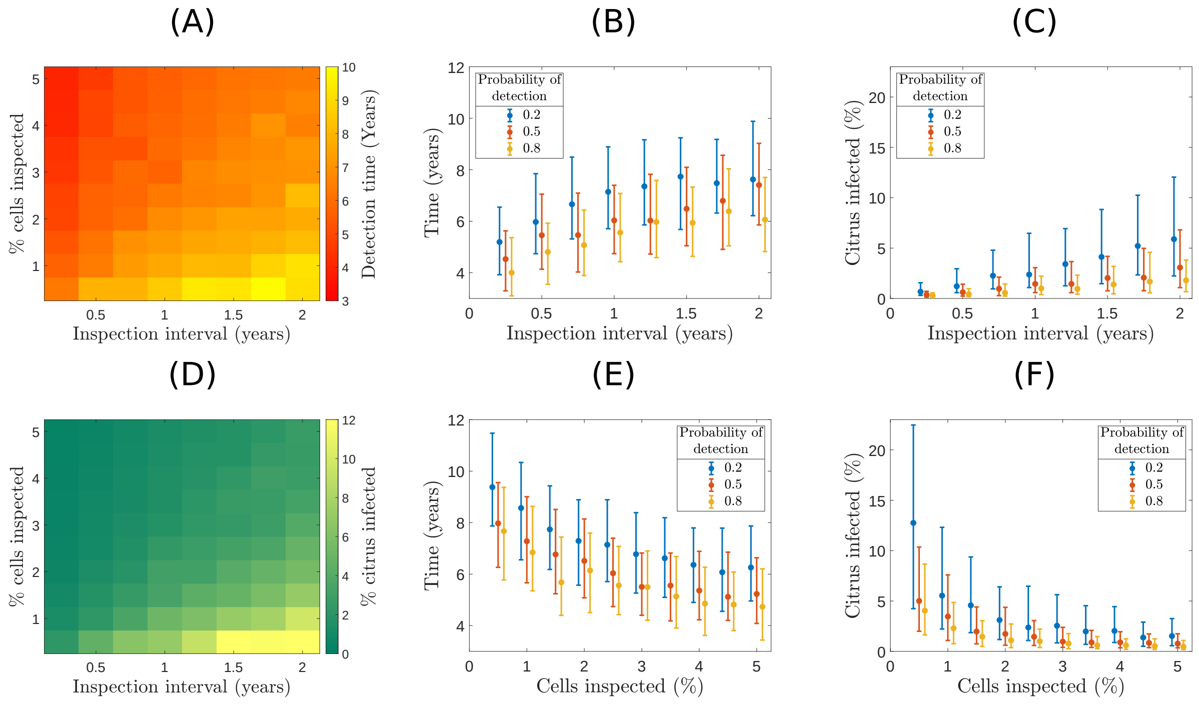


**Figure S17: Effectiveness of surveillance strategies when varying the inspection interval and amount of citrus that is inspected in Region B (Andalusia).** (A) Mean time of detection and (D) Mean percentage of total citrus infected at the time of detection. The number of units visited at each cell is $n_h = 5$ and the probability of detection is $p = 0.5$. (B,C,E,F) The median and inter-quartile range of (B,E) time until detection and (C,F) proportion of citrus that is infected. The number of samples inspected per cell is 5 and the probability of detection, inspection intervals and % of cells inspected are varied; (B,C) has fixed 5% of cells inspected at varying intervals, and (E,F) varies the % of cells inspected at a fixed 12 month inspection interval. The plots show responses for three values of the probability of detection, $p$.

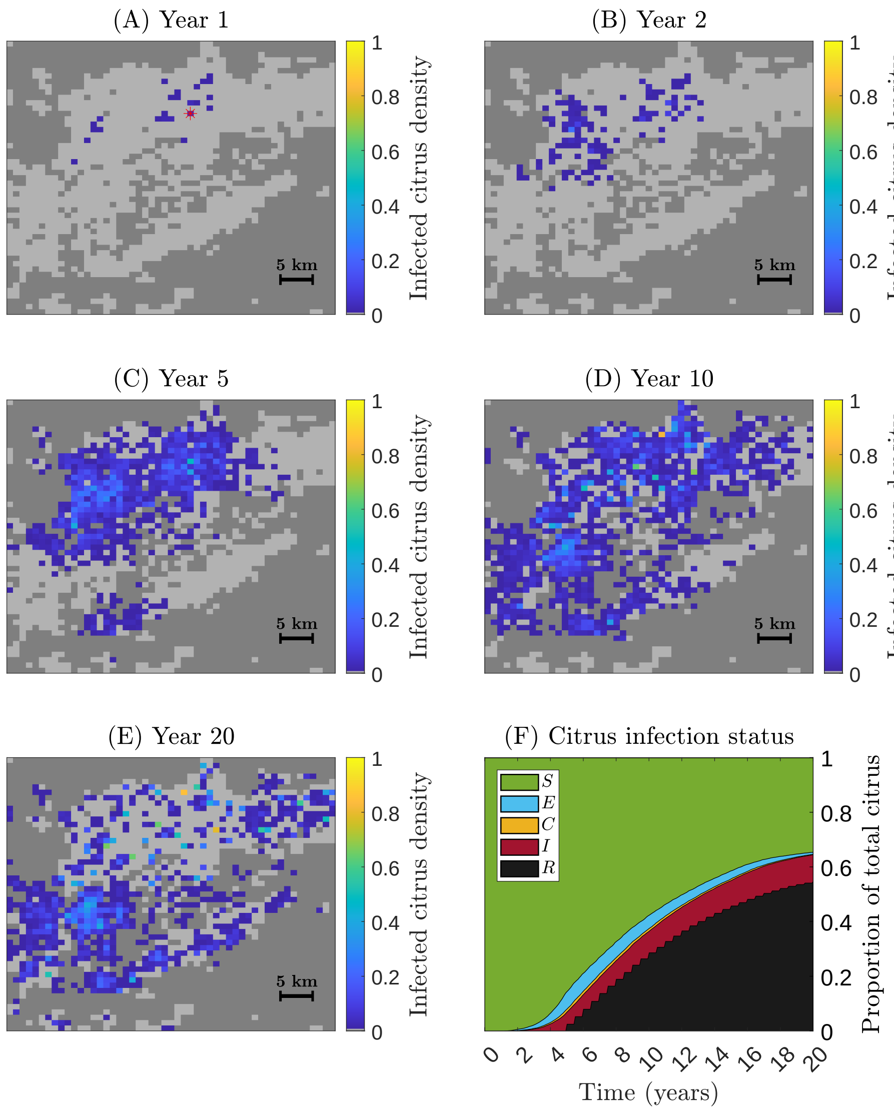


**Figure S18: An example from a single simulation of the spread of HLB over time in Region B (Andalusia) using baseline parameters for detection and control.** Maps show the density of infected citrus ($E + C + I$) per grid cell. In this example, the same initial conditions were used as Fig. S15 but included detection and control with the default parameterisation (Table 2), and so before the pathogen was detected, 5 host units within 1% of cells containing commercial citrus were inspected once per year, with a probability of detection $p = 0.5$. The disease was first detected after approximately 5 years. After that, all commercial units were inspected once every six months and any symptomatic citrus was rogued, with a 90% compliance rate and a 90% roguing probability. See also S3 Supporting Videos (Video 6).

### S3 Supporting Videos

Videos of simulation runs showing typical behaviour of our model for the following scenarios are archived at `doi:10.5281/zenodo.11222664`

#### S3.1 Spread in Region A (Valencia)

Spread of vector and pathogen without control.

- `Video1.avi` shows the spread of the vector (Fig. S10).
- `Video2.avi` shows the corresponding spread of the pathogen (Fig. 3 in main text).

Spread with control.

- `Video3.avi` shows spread of the pathogen (Fig. 6 in main text).

#### S3.2 Spread in Region B (Andalusia)

Spread of vector and pathogen without control.

- `Video4.avi` shows the spread of the vector (Fig. S14).
- `Video5.avi` shows the corresponding spread of the pathogen (Fig. S15).

Spread with control.

- `Video6.avi` shows spread of the pathogen (Fig. S18).
